# SARS-CoV-2 ORF6 disturbs nucleocytoplasmic trafficking to advance the viral replication

**DOI:** 10.1101/2021.02.24.432656

**Authors:** Yoichi Miyamoto, Yumi Itoh, Tatsuya Suzuki, Tomohisa Tanaka, Yusuke Sakai, Masaru Koido, Chiaki Hata, Cai-Xia Wang, Mayumi Otani, Kohji Moriishi, Taro Tachibana, Yoichiro Kamatani, Yoshihiro Yoneda, Toru Okamoto, Masahiro Oka

## Abstract

Severe acute respiratory syndrome coronavirus 2 (SARS-CoV-2) is the virus responsible for the coronavirus disease 2019 pandemic. ORF6 is known to antagonize the interferon signaling by inhibiting the nuclear translocation of STAT1. Here we show that ORF6 acts as a virulence factor through two distinct strategies. First, ORF6 directly interacts with STAT1 in an IFN-independent manner to inhibit its nuclear translocation. Second, ORF6 directly binds to importin α1, which is a nuclear transport factor encoded by *KPNA2*, leading to a significant suppression of importin α1-mediated nuclear transport. Furthermore, we found that *KPNA2* knockout enhances the viral replication, suggesting that importin α1 suppresses the viral propagation. Additionally, the analyses of gene expression data revealed that importin α1 levels decreased significantly in the lungs of older individuals. Taken together, SARS-CoV-2 ORF6 disrupts the nucleocytoplasmic trafficking to accelerate the viral replication, resulting in the disease progression, especially in older individuals.

## INTRODUCTION

Coronavirus disease 2019 (COVID-19) pandemic is caused by the severe acute respiratory syndrome coronavirus 2 (SARS-CoV-2), which is a single-strand RNA virus belonging to the *Coronaviridae* family ^1–3^. The genome of SARS-CoV-2 is approximately 29.7 kb long with short untranslated regions (UTR) at the 5’ and 3’ termini, and encodes nonstructural (nsp1-16), structural (spike [S], envelope [E], membrane [M], and nucleocapsid [N]), and accessory proteins (ORF3a, ORF3b, ORF6, ORF7a, ORF7b, ORF8, and ORF10) ^4, 5^.

Among them, ORF6 is a small protein of approximately 7 kDa, which consists of 61 amino acids and exhibits a 69% homology with the SARS-CoV ORF6, from which it differs due to a two amino acids deletion at the C-terminus ^6^. Several studies have recently shown that both the SARS-CoV and SARS-CoV-2 ORF6 proteins antagonize the host innate immune system via the Janus activated kinase 1 (JAK1)- and JAK2-signal transducers and activators of transcription (STAT) ^6–10^. STAT1 is a key mediator of cytokine-induced gene expression as it is activated by cytokines including type I and type II interferons (IFNs) ^11, 12^. Activation of JAKs associated with type I IFN receptor results in the tyrosine phosphorylation of STAT1 (PY-STAT1), leading to the formation of a STAT1-STAT2 heterodimer, while Type I interferon (IFN-α or -β) and type II interferon (IFN-γ) induce the formation of the PY-STAT1 homodimers. Both the hetero- and homo-dimer STAT1 complexes translocate to the nucleus to bind the IFN-stimulated response elements (ISRE) or IFN-γ-activated site (GAS) ^11, 12^. Previous studies indicated that ORF6 inhibits the nuclear transport of PY-STAT1 to suppress the primary interferon signaling ^7–10, 13^.

Constitutive and signal-dependent protein transports through the nuclear pore complexes (NPCs) embedded in the nuclear envelope are mediated by members of the importin (also known as karyopherin) superfamily ^14–17^. The process of protein import into the nucleus commonly involves the recognition of a classical nuclear localization signal (cNLS) by the importin α/β1 heterodimer ^18, 19^. Cargos containing the cNLS are recognized by the adaptor molecule importin α (also known as karyopherin α: KPNA). Following the cNLS-containing cargo/importin α/β1 trimeric complex entrance into the nucleus through the NPCs, the cargo is released from importin α by binding of GTP-bound small GTPase Ran (RanGTP) to importin β1 ^18–20^. After the complex dissociation, importin α is exported to the cytoplasm by cellular apoptosis susceptibility gene product (CAS, also known as CSE1L) with RanGTP, and the importin β1/RanGTP complex also returns to the cytoplasm, where it is reused for the next rounds of transport ^14, 20^.

Seven importin α proteins have been identified in humans, while six have been discovered in mice ^17, 20, 21^. Based on the sequence similarity, each importin α protein is assigned to one of three conserved subfamilies. In humans, clade 1 consists of importin α5 (encoded by the *KPNA1* gene), importin α6 (*KPNA5*), and importin α7 (*KPNA6*). Clade 2 consists of importin α1 (*KPNA2*) and importin α8 (*KPNA7*), and clade 3 consists of importin α3 (*KPNA4*) and importin α4 (*KPNA3*) ^20^. The members of each subfamily display the cargo specificity and are differentially expressed in different tissues and cell types ^20–23^. Differences in the usage of “importin α” or “KPNA” and the number of proteins in humans have often caused confusion. Therefore, in this study, we uniformly use the human nomenclature for “importin α” to refer to the protein and use the italic term “*KPNA*” to refer to the gene, while the normal term “KPNA” is described according to the way in which it is used in the citation.

It has already been demonstrated that the nuclear transport of the PY-STAT1 as a homodimer or a heterodimer with STAT2 is mediated by specific clade 1 subtypes of importin α such as importin α5/KPNA1 ^24–27^. According to the several studies on the viral proteins that inhibit the nuclear transport of PY-STAT1, SARS-CoV ORF6 has been reported to tether KPNA2, but not KPNA1, to ER to sequester importin β1 into endoplasmic reticulum (ER)/Golgi apparatus, resulting in the suppression of nuclear import of PY-STAT1 ^7^. Recently, SARS-CoV-2 ORF6 has also been shown to interact with KPNA2 ^10^ as well as both KPNA1 and KPNA2 ^9^, further supporting its interference with the nuclear transport of PY-STAT1. On the other hamd, SARS-CoV-2 ORF6 was shown to interact with the Nup98-RAE1 (NPC components) complex ^28^, and to achieve the nuclear exclusion of STAT1 by binding to the Nup98-RAE1 complex rather than binding to importin α proteins ^9^. However, the exact molecular mechanism of the effects of SARS-CoV-2 ORF6 on the nucleocytoplasmic trafficking remains largely unknown.

In this study, therefore, we characterized the function of SARS-CoV-2 ORF6 on the nucleocytoplasmic protein transport. First of all, we examined the subcellular localization of ORF6 protein in cells infected with distinct SARS-CoV-2 strains using originally established antibodies, and also demonstrated that ORF6 functions as a virulence factor for COVID-19 using a hamster model and a newly produced replicon system. Furthermore, we found that ORF6 directly binds to STAT1 to suppress the IFN-induced nuclear localization and the nuclear shuttling in the absence of IFN-stimulation. In addition, the direct binding of ORF6 to importin α1 significantly reduces the cNLS-cargo transport, indicating that ORF6 influences importin α1 independently of STAT1. We also found that the viral replication of SARS-CoV-2 is enhanced in *KPNA2* knockout cells, suggesting that importin α1 acts for the suppression of viral propagation. Lastly, the analyses of datasets from Genotype-Tissue Expression (GTEx) project indicated that the expression of importin α1 significantly decreases in the lungs of older individuals, suggesting that the importin α1 levels might be related to the expansion of the COVID-19 illness. Thus, our results suggest that ORF6 plays a critical role in SARS-CoV-2 replication by disrupting the importin α-mediated nucleocytoplasmic protein trafficking.

## RESULTS

### ORF6 contributes to viral RNA replication and pathogenicity *in vivo*

To assess the ORF6 expression in SARS-CoV-2 infected cells, we first established an ORF6 specific antibody, and examined the protein in cells infected with different viral strains which were obtained from the National Institute of Infectious Diseases (NIID) in Japan, Hong Kong (HK)/VM20001061, USA-CA2, Germany/BavPat1, New York (NY)-PV09197, NY-PV08410, and NY-PV08449 (Fig. 1A). Indirect immunofluorescence analysis indicated that ORF6 was localized in the cytoplasm of the SARS-CoV-2 infected VeroE6/TMPRSS2 cells (Fig. 1B), which is consistent with the previous observation in SARS-CoV infected Vero E6 cells ^29^.

**Figure 1.**
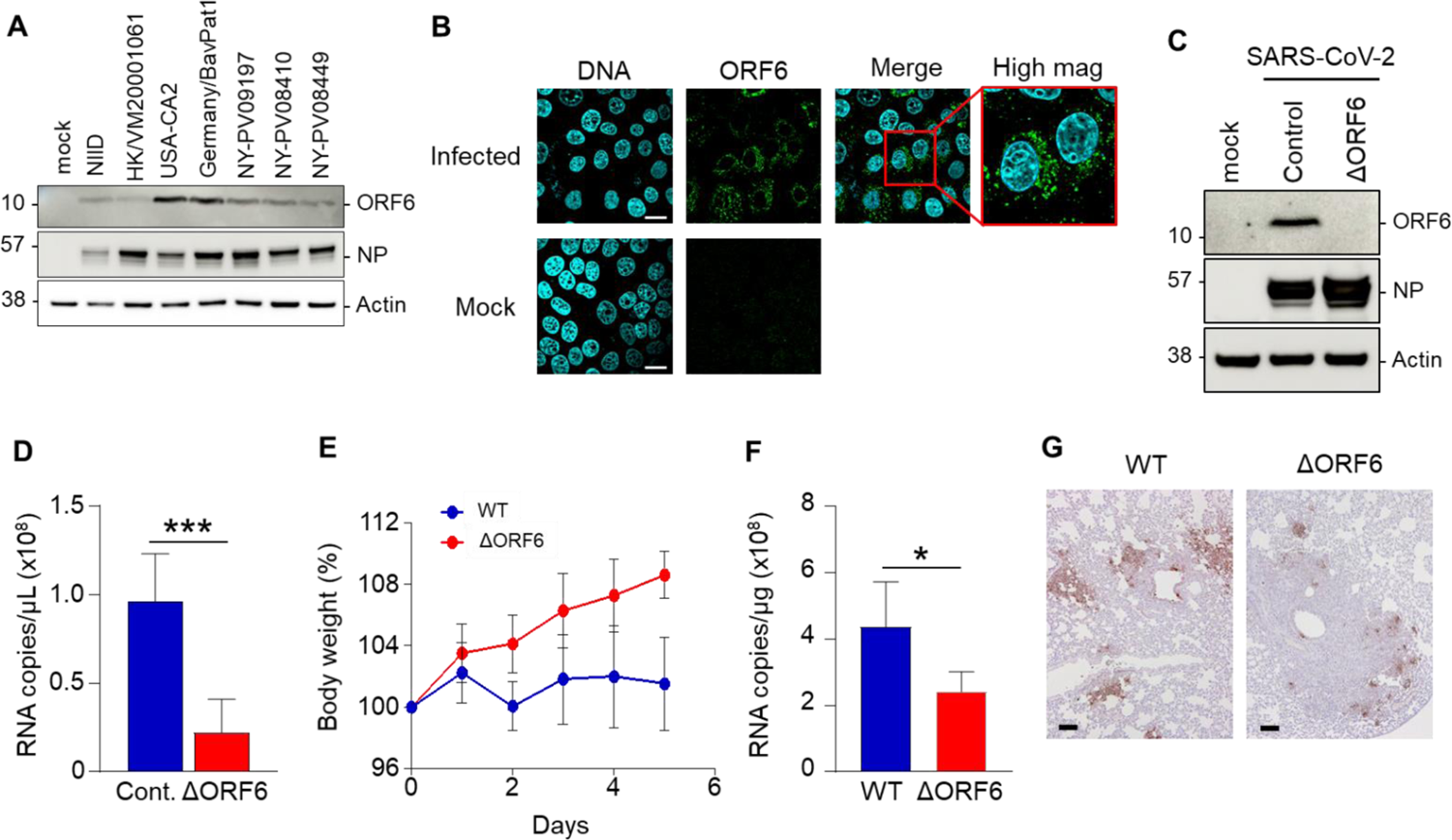
Mutated viruses reveal the virulence of ORF6 in SARS-CoV-2 replication. **A.** Identification of ORF6 in different SARS-CoV-2 strains by a newly established antibody. VeroE6/TMPRSS2 cells were infected with several SARS-CoV-2 strains, and cell lysates were collected for detection of the ORF6 protein using western blotting. NIID: 2019-nCoV/Japan/TY/WK-521/2020 strain was isolated at the National Institute of Infectious Diseases. The nucleoprotein (NP) and Actin were used as an infection control and internal control, respectively. **B.** Indirect immunofluorescence of ORF6 in VeroE6/TMPRSS2 cells infected with or without (sham) SARS-CoV-2 from NIID. The squares with a red line show a magnified marge image. Scale bars: 20 μm. **C.** VeroE6/TMPRSS2 cells were infected with SARS-CoV-2 WT (Nluc-2A-ORF6) or ΔORF6 and the cell lysates were collected at 24 h post infection. Cell lysates were subjected to western blotting, and then detected the proteins using specific antibodies for ORF6, NP, or Actin. **D.** Huh7-ACE2 cells were infected with SARS-CoV-2 WT (Nluc-2A-ORF6) or ΔORF6 and supernatants were collected at 24 h post infection. Viral RNA in the supernatants was quantified using qRT-PCR. ***P < 0.001, two-tailed Student’s t-test. **E.** Percent body weight changes were calculated for all hamsters infected with SARS-CoV-2 WT or ΔORF6. Data are mean ± SD from four independent animals. **F.** Viral RNA in lung homogenates from hamsters was quantified using qRT-PCR. *P < 0.05, two-tailed Student’s t-test. **G.** Immunohistochemistry of SARS-CoV-2 antigen (NP protein) in lung lobes of hamster infected with SARS-CoV-2 WT or ΔORF6, respectively. Scale bars: 100 μm.

Next, we tried to know the roles of ORF6 in the viral life cycle of SARS-CoV-2. For this, the viral genome encoding ORF6 was replaced by that of NanoLuc, and a SARS-CoV-2 variant that does not express ORF6 (SARS-CoV-2/ΔORF6) was generated using the circular polymerase extension reaction (CPER). As a parental virus, a recombinant virus expressing ORF6 fused with NanoLuc (NLuc) and Porcine teschovirus 2A peptide (SARS-CoV-2/NLuc2AORF6, Fig. S1) was generated. The deletion of ORF6 in SARS-CoV-2/ΔORF6 was confirmed by western blotting (Fig. 1C). To assess the viral growth in Huh7-ACE2 cells, these recombinant viruses were inoculated at a MOI = 0.1, and then the viral RNA was determined from the culture supernatant. The viral RNA in the supernatant of cells infected with SARS-CoV-2/ΔORF6 was significantly decreased (Fig. 1D).

Next, to evaluate the function of ORF6 *in vivo*, the parental virus (WT SARS-CoV-2) or SARS-CoV-2/ΔORF6 was intranasally inoculated in 4-week-old hamsters. Hamsters infected with SARS-CoV-2/ΔORF6 showed significant weight gain 5 days post-infection, whereas those infected with the WT virus showed no significant change in the body weight (Fig. 1E). In addition, 5 days post-infection, the viral RNA was significantly reduced in the lung cells infected with SARS-CoV-2/ΔORF6 compared to those infected with the WT virus (Fig. 1F). Immunohistological analyses revealed that the viral nucleoprotein (NP) was expressed at lower levels in the lung cells infected with SARS-CoV-2/ΔORF6 than in those infected with the WT virus (Fig. 1G). These data suggest that ORF6 is involved in the viral replication and pathogenesis of COVID-19 *in vivo*.

### ORF6 inhibits the nuclear localization of STAT1 following the IFN stimulation

Several studies have already shown that ORF6 inhibits the nuclear localization of STAT1 in response to type-I IFN (IFN-α or -β) stimulation ^7–10, 13^. Here, we attempted to verify the inhibitory effect of SARS-CoV-2 ORF6 on a type-II IFN (IFN-γ)-activated STAT1. The AcGFP fused ORF6 was transfected into HeLa cells, and the subcellular localization of PY-STAT1 was observed when the cells were stimulated with either IFN-β or IFN-γ. While the PY-STAT1 was localized in the nucleus of AcGFP-transfected cells in response to each treatment, the distribution was markedly shifted to the cytoplasm in the AcGFP-ORF6-transfected cells (Fig. 2A, B). The fluorescence intensity ratio of the nucleus against the whole cells further supported the significant nuclear exclusion of the PY-STAT1 in the AcGFP-ORF6-transfected cells when stimulated by IFN-γ (Fig. 2C). Next, in order to confirm that the PY-STAT1 is excluded from the nucleus by ORF6, we evaluated the expression of PY-STAT1 downstream genes using quantitative RT-PCR (qRT-PCR). In comparison to the AcGFP-transfected control cells, the AcGFP-ORF6-transfected cells showed significant down-regulation of IFN-γ-inducible protein 10 (*IP-10*) ^30^ mRNA 6 h post-transfection or later upon stimulation (Fig. 2D). To clarify whether the nuclear exclusion of PY-STAT1 by ORF6 affects the expression of the IFN-stimulated response element (ISRE)-containing gene, a luciferase assay was performed in Huh7 cells. The cells were transfected with a luciferase reporter plasmid including ISRE together with AcGFP or AcGFP-ORF6. We observed a significant repression of the relative luciferase values in the AcGFP-ORF6-transfected cells compared to that of AcGFP-transfected cells (Fig. 2E). Finally, we confirmed that the relative ISRE luciferase value was significantly suppressed when the SARS-CoV-2 infected cells were stimulated by IFN-γ (Fig. 2F). These results indicate that SARS-CoV-2 ORF6 suppresses the nuclear translocation of PY-STAT1 to inhibit the activation of STAT1-downstream genes.

**Figure 2.**
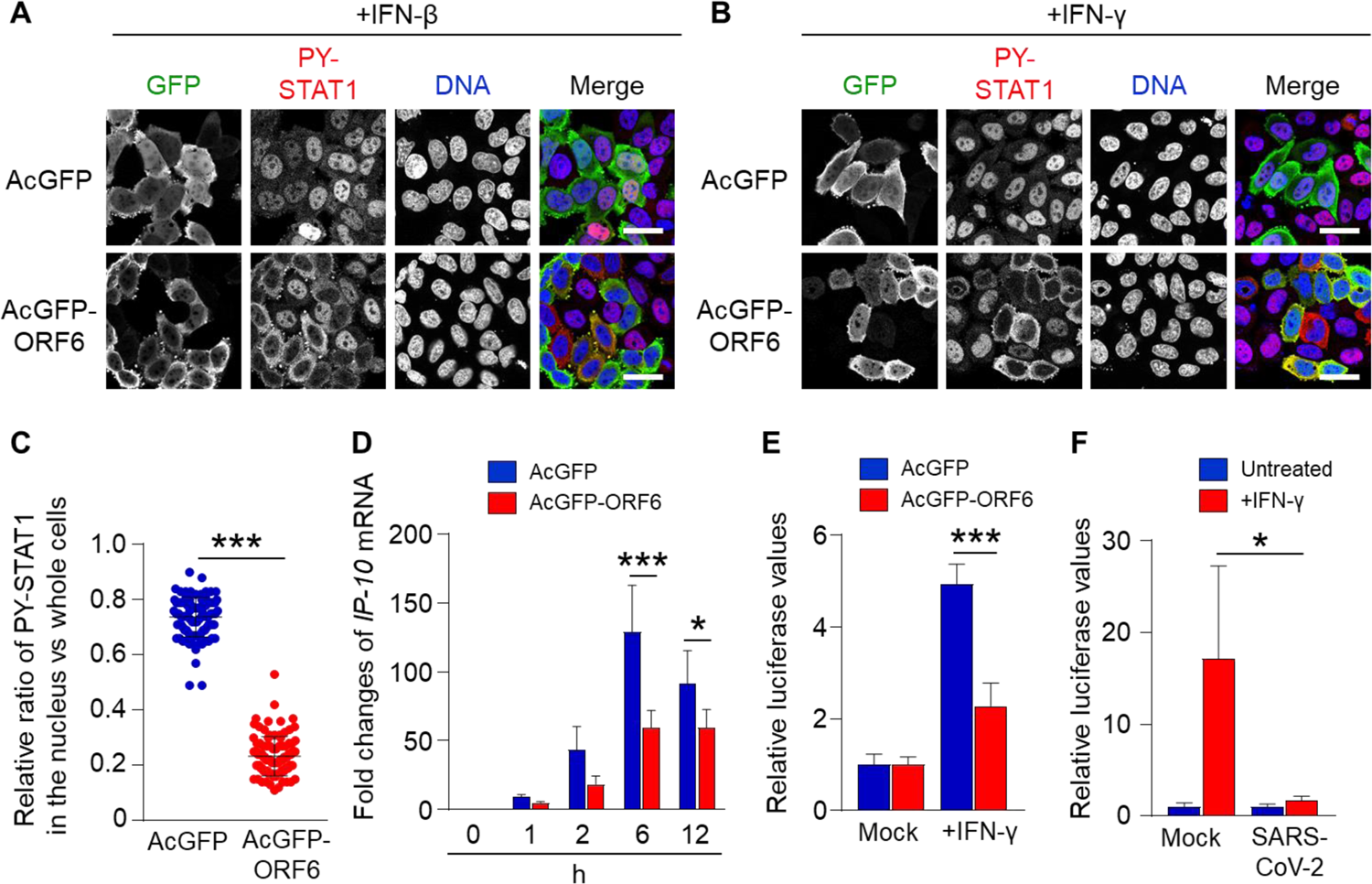
Inhibition of nuclear localization of PY-STAT1 by ORF6. **A-B.** Immunofluorescence of phosphorylated STAT1 (PY-STAT1) in HeLa cells transfected with AcGFP or AcGFP-ORF6 following INF-β (**A**) or IFN-γ (**B**). AcGFP (green) and PY-STAT1 (red) were identified using specific antibodies. DAPI staining (blue) was used for DNA staining. Scale bars: 30 μm. **C.** The graph represents the relative fluorescence values of PY-STAT1 in the nucleus compared to those of the whole cells (shown in **B**). Signal intensities of total 100 different nuclei from two independent experiments. ***P < 0.001, two-tailed Student’s t-test. **D.** qRT-PCR analysis of *IP-10* mRNA in AcGFP and AcGFP-ORF6-transfected HeLa cells at the described time points (n = 4 each). *P < 0.05, ***P < 0.001, two-way ANOVA. **E.** Relative luciferase values of ISRE-TA-Luc in AcGFP and AcGFP-ORF6-transfected Huh7 cells upon IFN-γ stimulation. ***P < 0.001, one-way ANOVA. **F.** Relative luciferase values in VeroE6/TMPRSS2 cells infected with SARS-CoV-2 (NIID strain). *P < 0.05, one-way ANOVA.

### The C-terminal region of ORF6 aids the acceleration of viral replication

Previously, the C-terminal region of ORF6 was shown to be involved in the inhibition of IFN response ^8^. Hence, in line with the previous findings, we validated the effects of ORF6 C-terminal mutations on the nuclear translocation of PY-STAT1. As a result, we found that the ORF6 mutant with alanine replacing amino acids 49 to 52 (referred to as ORF6-M1) retained the inhibitory effect over PY-STAT1, while the other mutants with alanine replacing amino acids 53 to 55 (ORF6-M2) and 56 to 61 (ORF6-M3) did not retain their inhibitory functions (Fig. 3A-C).

**Figure 3.**
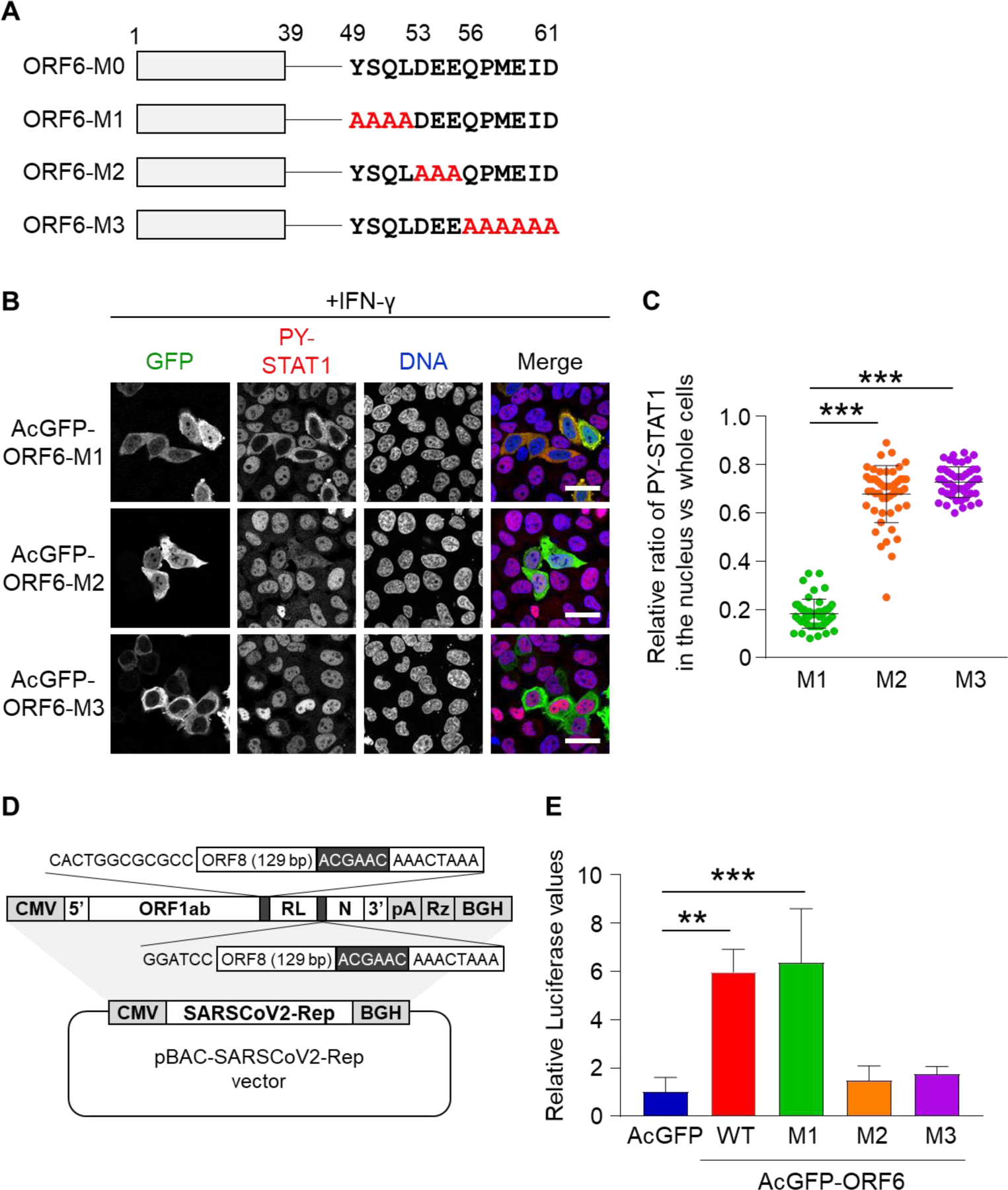
The C-terminal region of ORF6 contributes to viral RNA replication. **A.** Schematic representation of ORF6 wild type (WT) and its alanine substitution mutations within the C-terminus. ORF6-M0: wild type ORF6; ORF6-M1: ORF6 with amino acids 49-52 substituted for alanine; ORF6-M2: ORF6 with amino acids 53-55 substituted for alanine; ORF6-M3: ORF6 with amino acids 56-61 substituted for alanine. **B.** Immunofluorescence of PY-STAT1 in HeLa cells transfected with the AcGFP-ORF6 mutants following IFN-γ stimulation. DAPI was used to stain the DNA. Scale bars: 30 μm. **C.** The graph represents the relative fluorescence values in the nucleus compared to those of the whole cells in **B**. Signal intensities of total 50 nuclei from two independent experiments were measured. ***P < 0.001, one-way ANOVA. **D.** Schematic representation of SARS-CoV-2 replicon DNA, pBAC-SCoV2-Rep. The genetic structure of the SARS-CoV-2 replicon is shown at the top of the panel. The dark shaded box indicates the core sequence of transcription regulating sequence. CMV, cytomegalovirus promoter; RL, Renilla luciferase gene; pA, a synthetic poly(A) tail; Rz, hepatitis delta virus ribozyme; BGH, bovine growth hormone polyadenylation sequence. **E.** Relative luciferase values for each ORF6 mutant in replicon (n=3). **P < 0.01, ***P < 0.001, one-way ANOVA.

To further assess the function of ORF6 in viral propagation, we established a replicon system in which we can evaluate the viral replication process by detecting the Renilla luciferase (RLuc) (Fig. 3D). The replicon plasmid was transfected together with AcGFP or AcGFP-ORF6 in Huh7 cells and the RLuc values were analyzed 24 h post-transfection. We found that the expression of WT ORF6 and ORF6-M1, but not ORF6-M2 and ORF6-M3, significantly enhanced the viral replication (Fig. 3E), consistent with the effects on STAT1 nuclear localization. These results suggest that the C-terminal regions (amino acids 53 to 61) of ORF6 plays an important role to inhibit the interferon signaling by disrupting the nuclear localization of PY-STAT1, resulting in the enhancement of the viral replication of SARS-CoV-2.

### ORF6 directly binds to STAT1 in an IFN-independent manner

Next, in order to characterize the interplay between ORF6 and STAT1 in more detail, we analyzed the subcellular distribution of Flag-tagged STAT1 in AcGFP-ORF6-transfected HeLa cells. In addition to the WT ORF6, the in-frame 9 amino acids deletion mutant (loss of amino acids 22 to 30; referred to as ORF6Δ9) which were reported in the previous works ^31, 32^ was also examined. Under non-stimulated conditions, Flag-STAT1 was mainly localized in the cytoplasm in either AcGFP-, AcGFP-ORF6- or AcGFP-ORF6Δ9-transfected cells (Fig. 4A, B). Upon IFN-γ stimulation, Flag-STAT1 was distributed into the nucleus in the AcGFP-transfected cells. In contrast, it was retained in the cytoplasm in AcGFP-ORF6 WT- and Δ9-transfected cells (Fig. 4C, D). Interestingly, statistical analysis revealed that even under non-stimulated conditions, the cytoplasmic intensities of Flag-STAT1 were significantly higher in the AcGFP-ORF6 WT- or Δ9-transfected cells compared to those observed in the AcGFP-transfected control cells (Fig. 4B), suggesting that STAT1 may shuttle between the nucleus and the cytoplasm in the absence of IFN stimulation and be trapped in the cytoplasm by ORF6.

**Figure 4.**
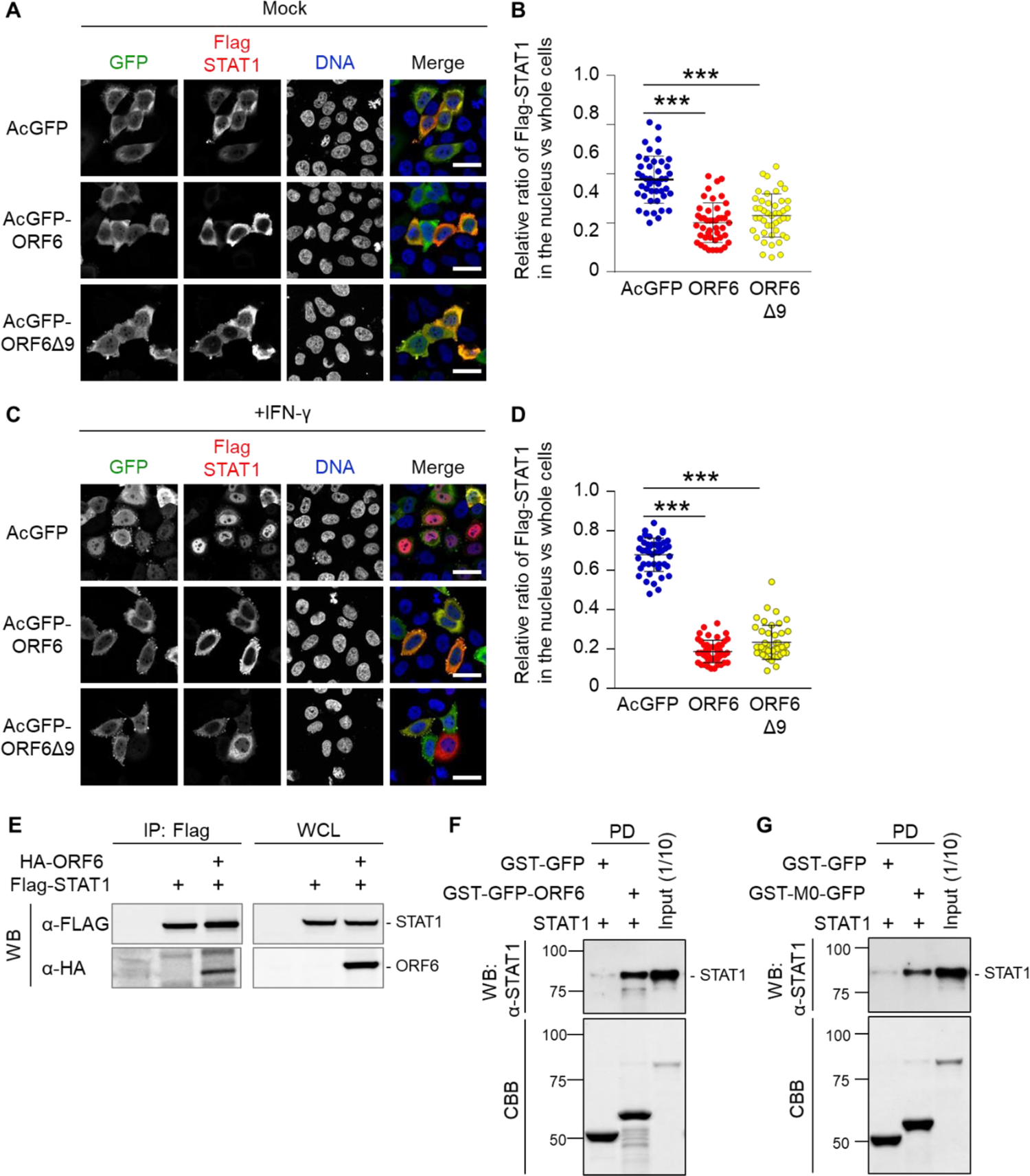
ORF6 directly binds to STAT1. **A.** Immunofluorescence of Flag-STAT1 in HeLa cells transfected with AcGFP, AcGFP-ORF6 WT or AcGFP-ORF6Δ9 (anti-GFP or anti-Flag antibodies were used). DAPI was used to stain the DNA. Scale bars: 30 μm. **B.** The graph represents the relative fluorescence values of Flag-STAT1 in the nucleus compared to those of the whole cells in **A**. Signal intensities from total 45 nuclei from two independent experiments. ***P < 0.001, one-way ANOVA. **C.** Immunofluorescence of Flag-STAT1 in HeLa cells transfected with AcGFP, AcGFP-ORF6 WT or AcGFP-ORF6Δ9 following IFN-γ stimulation. Anti-GFP or anti-Flag antibodies were used for detection. DAPI was used to stain the DNA. Scale bars: 30 μm. **D.** The graph represents the relative fluorescence values of Flag-STAT1 in the nucleus compared to those of the whole cells in **C**. Signal intensities from total 45 nuclei from two independent experiments were measured. ***P < 0.001, one-way ANOVA. **E.** Immunoprecipitation of ORF6 was used to detect the binding with STAT1. HEK293 cells were transiently transfected with HA fused AcGFP-ORF6 (HA-ORF6) and Flag-STAT1 for 24 h. Cell lysates were subjected to immunoprecipitation using an anti-Flag antibody. Following the immunoprecipitation, the samples were western blotted with the anti-Flag antibody or the anti-HA antibody, respectively. **F.** GST-GFP and GST-GFP-ORF6 immobilized on glutathione Sepharose beads were incubated with the bacterially purified STAT1 recombinant protein for 1 h. The bottom panel represents the proteins bound to the beads and stained with Coomassie Brilliant Blue (CBB). Input is 1/10th of the amount of STAT1 used for the reaction. **G.** GST-GFP or GST-GFP fused with the C-terminal peptide of ORF6 wild type (49-61 amino acids; GST-M0-GFP) immobilized on glutathione Sepharose beads was incubated with the STAT1 recombinant protein for 1 h. The bottom panel represents the proteins bound to the beads and stained with CBB. Input is 1/10th of the amount of STAT1 used for the reaction.

Therefore, to address the possibility of the interaction between ORF6 and non-activated STAT1, we first performed an immunoprecipitation assay using AcGFP-ORF6 (HA-ORF6)- and Flag-STAT1-transfected cells. As a result, we found that HA-ORF6 was precipitated with Flag-STAT1 in the absence of IFN stimulation (Fig. 4E). Next, to know whether ORF6 directly binds to STAT1, bacterially purified recombinant STAT1 protein was incubated with either recombinant GST-GFP or recombinant GST-GFP fused ORF6 full-length protein (GST-GFP-ORF6). Fig. 4F showed that STAT1 was directly bound to GST-GFP-ORF6. To address whether the binding of ORF6 to STAT1 is mediated by the amino acids 49-61 in the C-terminal region (referred to as M0 in this study, see Fig. 3A), we produced a recombinant protein consisting of the 13 amino acids (M0) which was fused to GST and GFP (GST-M0-GFP). As shown in Fig. 4G, the pull-down assay clearly revealed that STAT1 was bound to the GST-M0-GFP protein. Taken together, we conclude that ORF6 directly binds to STAT1 through the C-terminus in the absence of IFN stimulation, meaning that ORF6 binds to the resting state of STAT1 in the cytoplasm.

### ORF6 alters the subcellular distribution of importin α proteins

Previous studies showed that the SARS-CoV- and SARS-CoV-2-associated inhibition of nuclear translocation of STAT1 is accomplished through the interaction between ORF6 and importin α1/KPNA2 ^7, 10^. However, as described above, ORF6 binds directly to STAT1 in the absence of IFN stimulation in the cytoplasm, which raises a possibility that the binding of ORF6 to importin α proteins might not be required to inhibit the nuclear accumulation of STAT1. Therefore, we attempted to examine the interplay between the ORF6 and importin α proteins. Since ORF6 has been shown to alter the distribution of importin α1/KPNA2 and/or importin α5/KPNA1 ^7, 9^, we first validated these findings concerning all human importin α subtypes. Consistent with the previous reports, in AcGFP-transfected cells (control), the overexpressed Flag-importin α proteins were mainly localized in the nucleus (Fig. 5A, B, Fig. S2). In contrast, in AcGFP-ORF6-transfected cells, the localization of Flag-importin α1, α3, α4, α6, and α8 remarkably changed to the cytoplasm, while Flag-importin α5 and α7 were still mostly localized in the nucleus (Fig. 5A, B, Fig. S2). The analysis of nuclear fluorescence intensities ratio clearly showed that in AcGFP-ORF6-transfected cells, the nuclear distribution of Flag-importin α1 was drastically shifted to the cytoplasm, while the Flag-importin α5 was mainly retained in the nucleus (Fig. 5C). These data indicate that ORF6 has distinct effects on each importin α subtype.

**Figure 5.**
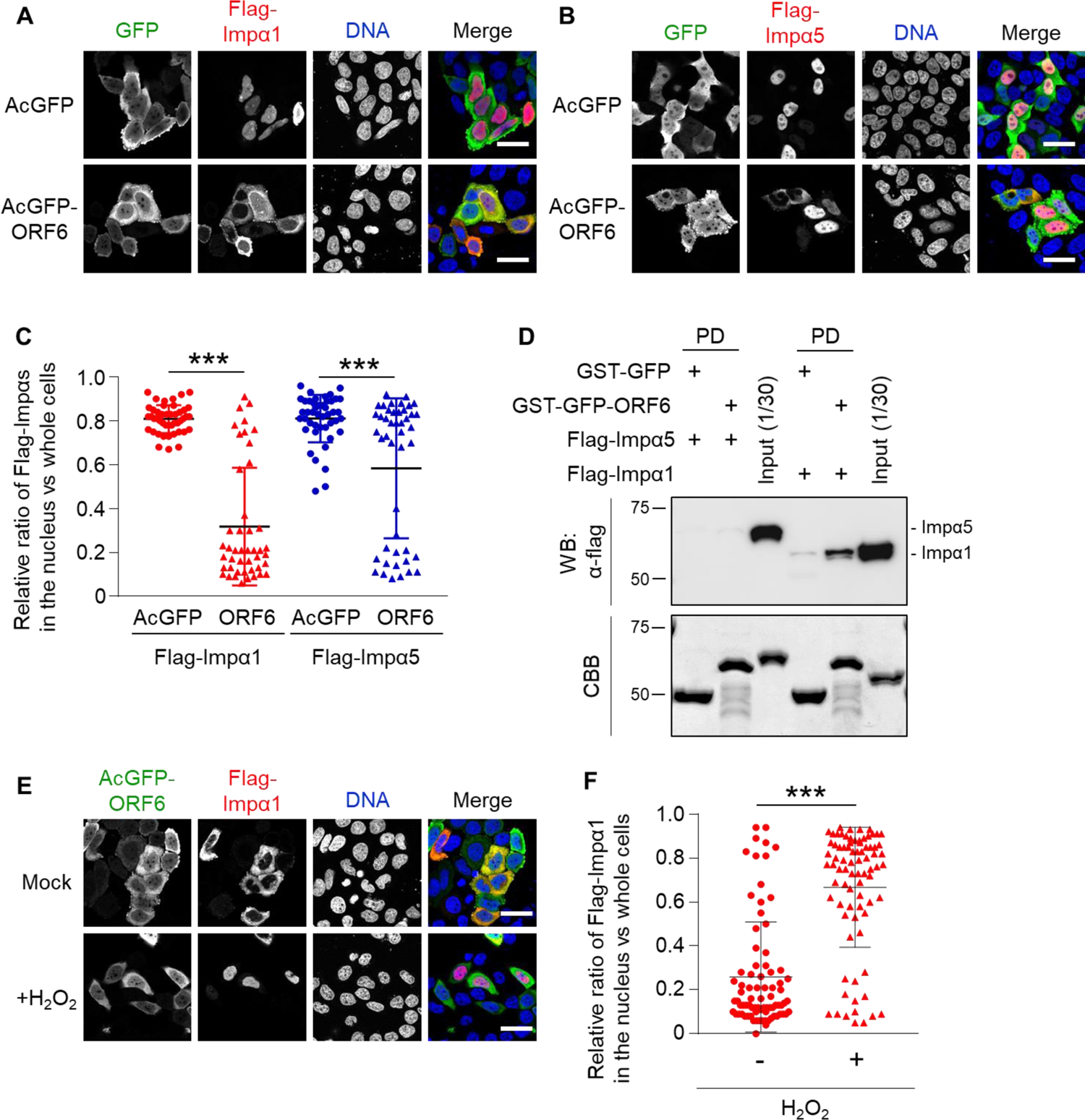
ORF6 affects the subcellular localization of importin α proteins. **A-B.** Immunofluorescence of Flag-importin α1 (Flag-Impα1) and Flag-importin α5 (Flag-Impα5) in HeLa cells transfected with AcGFP or AcGFP-ORF6. Anti-GFP or anti-Flag antibodies were used for detection. DAPI was used to stain the DNA. Scale bars: 30 μm. **C.** The graph represents the relative fluorescence values of the nucleus compared to those of the whole cells in **A** and **B**. Signal intensities of total 50 nuclei from two independent experiments were measured. ***P < 0.001, two-tailed Student’s t-test. **D.** GST-GFP and GST-GFP-ORF6 were immobilized on glutathione Sepharose beads and incubated with bacterially purified Flag-importin α1 (Flag-Impα1) or Flag-importin α5 (Flag-Impα5) for 1 h. The importin α proteins were detected using an anti-Flag antibody. The bottom panel represents the proteins bound to the beads and stained with CBB. Inputs are 1/30th of the amount of each importin α that was used for the reaction. **E.** Immunofluorescence of Flag-importin α1 (Flag-Impα1) in HeLa cells transfected with AcGFP-ORF6 with or without hydrogen peroxide (200 μM H_2_O_2_) for 30 min. Anti-GFP or anti-Flag antibodies were used for detection. DAPI was used to stain the DNA. Scale bars: 30 μm. **F.** The graph represents the relative fluorescence values of the nucleus compared to those of the whole cells in **E**. Signal intensities of total 80 nuclei from two independent experiments were measured. ***P < 0.001, two-tailed Student’s t-test.

### Importin α1 shuttles between the nucleus and the cytoplasm in the presence of ORF6

Next, we examined whether ORF6 directly binds to importin α proteins. Purified recombinant GST-GFP-ORF6 was incubated with Flag-importin α1 or Flag-importin α5, respectively, and then the GST-proteins were pulled-down using glutathione Sepharose beads to know whether the Flag-importin α proteins were co-precipitated or not. Using western blotting, we identified that Flag-importin α1, but not Flag-importin α5, mainly binds to ORF6 (Fig. 5D).

Direct binding of ORF6 to importin α1 suggest a possibility that ORF6 may inhibit the mobility of importin α1 to tether it in the cytoplasm, as reported previously with SARS-CoV ORF6 ^7^. Therefore, we next tried to know whether importin α1 moves from the cytoplasm to the nucleus even in the presence of ORF6. For this, we focused on the characteristic feature of importin α1 that it accumulates in the nucleus in response to cellular stresses such as oxidative stress ^33, 34^. That is, it has been clearly demonstrated that while in unstressed cells importin α1 shuttles between the nucleus and the cytoplasm, in stressed cells importin α1 accumulates in the nucleus due to the inhibition of RanGTP-dependent nuclear export of importin α by a collapse in the RanGTP gradient ^33, 34^. Therefore, we speculated that if importin α1 cannot move from the cytoplasm to the nucleus by ORF6, we cannot observe its nuclear accumulation under stress conditions. Hence, HeLa cells were transfected with AcGFP-ORF6 and Flag-importin α1 and treated with 200 μM of H_2_O_2_ for 1 h. Unexpectedly, we found that under the oxidative stress conditions, Flag-importin α1 was localized into the nucleus in the ORF6-transfected cells (Fig. 5E, F). The same phenomenon was also observed for endogenous importin α1 (Fig. S3A-C). Thus, these results mean that although importin α1 seems to be tethered in the cytoplasm in the ORF6-transfected cells, it retains its original capacity to move between the nucleus and the cytoplasm even in the ORF6-transfected cells. Taken together with the results that the subcellular localization of neither importin β1 nor CAS, the export factor for importin α, was dramatically altered in the ORF6-transfected cells (Fig. S3D, E), it is most likely that the nuclear exclusion of STAT1 by ORF6 is not due to the cytoplasmic tethering of importin α1.

### ORF6 negatively regulates the importin α/β1 pathway

As described above, since we found that ORF6 directly binds to importin α1, we next tried to know whether ORF6 directly affects the importin α1-mediated nuclear transport pathway or not. For this, HeLa cells were transfected with mCherry fused SV40T-NLS (mCherry-NLS) together with AcGFP or AcGFP-ORF6, and then the nuclear intensities of the fluorescent substrate were measured. Although the majority of mCherry-NLS was observed in the nucleus of AcGFP-ORF6-transfected cells, its cytoplasmic proportion was significantly increased compared to that observed in the AcGFP-transfected control cells (Fig. 6A, B).

**Figure 6.**
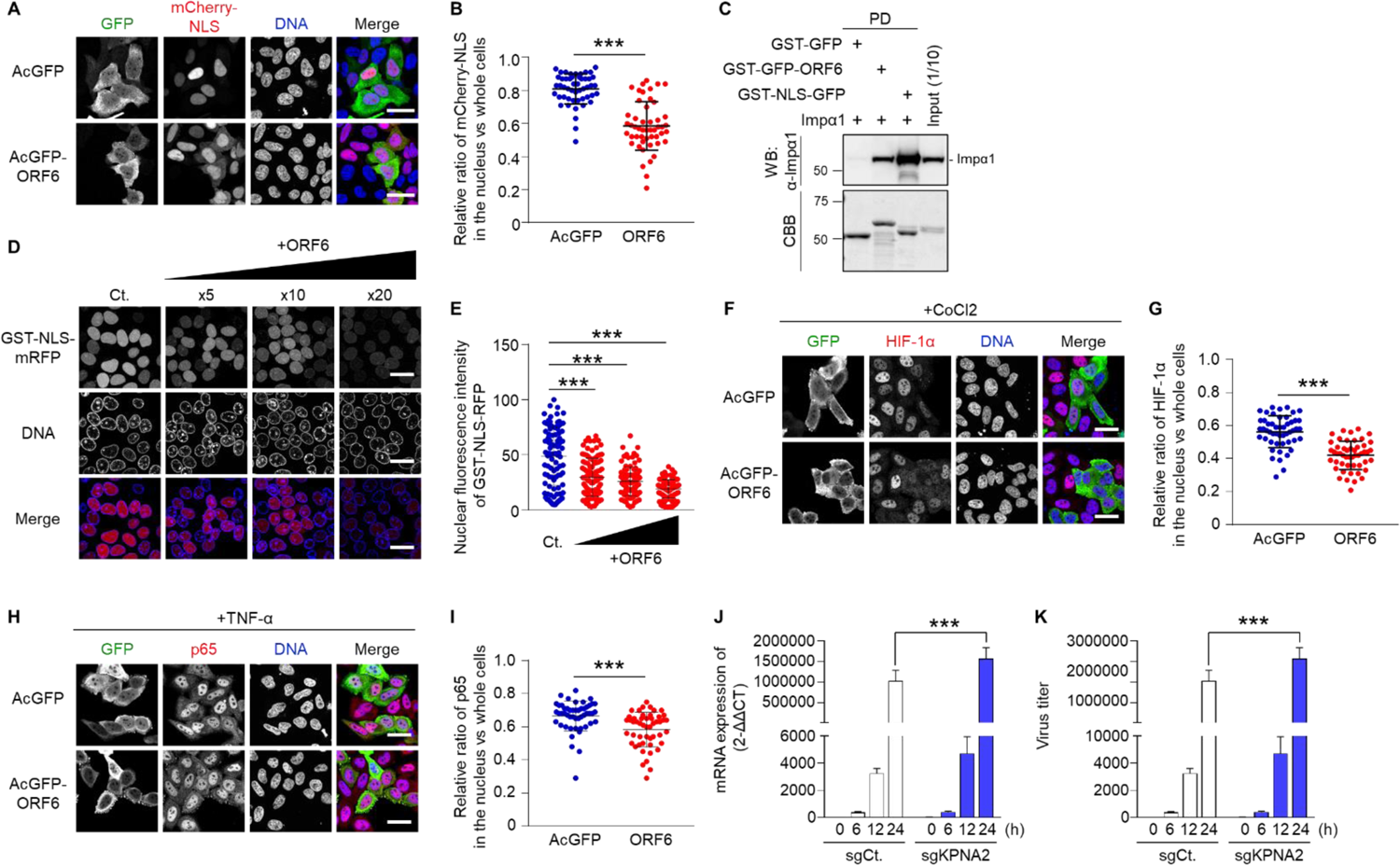
ORF6 negatively regulates the importin α/β1 pathway. **A.** Subcellular localization of mCherry-NLS in HeLa cells transfected with AcGFP or AcGFP-ORF6. DAPI was used to stain the DNA. Scale bars: 30 μm. **B.** The graph represents the relative fluorescence values of the nucleus compared to those of the whole cells in **A**. Signal intensities of total 50 nuclei from two independent experiments were measured. ***P < 0.001, two-tailed Student’s t-test. **C.** GST-GFP, GST-GFP-ORF6 or GST-NLS-GFP were immobilized on glutathione Sepharose beads and incubated with importin α1 (Impα1) for 1 h. The bottom panel represents the proteins bound to the beads and stained with CBB. Inputs are 1/10th of the amount of each importin α that was used for the reaction. **D.** An *in vitro* semi-intact nuclear transport assay was performed to measure the nuclear import of GST-NLS-mRFP in the presence of AcGFP-ORF6. Digitonin-permeabilized HeLa cells were incubated with GST-NLS-mRFP, importin α1, importin β1, RanGDP, p10/NTF2, GTP, and ATP regeneration system. The reaction mixture was added 5×, 10×, or 20× concentration of AcGFP-ORF6 compared to that of the NLS-substrate. After incubation for 30 min, the mRFP signals were detected using a fluorescence microscope. DAPI was used to stain the DNA. Scale bars: 30 μm. **E.** The graph represents the nuclear fluorescence values of GST-NLS-mRFP in **D**. Signal intensities of total 100 nuclei were measured and the statistically analyzed using a one-way ANOVA (***P < 0.001). **F**. Immunofluorescence of HIF-1α in HeLa cells transfected with AcGFP or AcGFP-ORF6 following CoCl2 treatment. Anti-GFP or anti-Flag antibodies were used for detection. DAPI was used to stain the DNA. Scale bars: 30 μm. **G**. The graph represents the relative fluorescence values of the nucleus compared to those of the entire cells in **F**. Signal intensities of total 50 nuclei from two independent experiments were measured. ***P < 0.001, two-tailed Student’s t-test. **H**. Immunofluorescence of NF-κB p65 in HeLa cells transfected with AcGFP or AcGFP-ORF6 following TNF-α stimulation. Anti-GFP or anti-Flag antibodies were used for detection. DAPI was used to stain the DNA. Scale bars: 30 μm. **I**. The graph represents the relative fluorescence values of the nucleus compared to whole cells in **H**. Signal intensities of total 50 nuclei from two independent experiments were measured. ***P < 0.001, two-tailed Student’s t-test. **J-K.** Huh7 cells expressing the ACE2 receptor (Huh7-ACE2) introduced with sgControl (sgCtl) or sgKPNA2 were infected with SARS-CoV-2 and supernatants were collected at 0, 6, 12, and 24 h. Intracellular viral RNA was quantified using qRT-PCR (**J**) while the viral titers (**K**) were quantified using plaque forming assay. Statistical significance was determined using a two-way ANOVA (***P < 0.001).

Next, we compared the binding ability of importin α1 to the cNLS-containing cargo (GST-NLS-GFP) and ORF6 (GST-GFP-ORF6). The GST pull-down assay showed that importin α1 more efficiently interacted with cNLS than ORF6 (Fig. 6C). To assess the inhibitory effects of ORF6 on the importin α/β1-mediated nuclear transport of GST-NLS-mRFP, a digitonin-permeabilized semi-intact nuclear transport assay was performed. As a result, we observed that the addition of excess amounts of ORF6 significantly inhibited the nuclear translocation of the cNLS-cargo (Fig. 6D, E), suggesting that ORF6 affects the importin α/β1 pathway, when it exists in large quantities in cells.

To further investigate the effects of ORF6 on other important signaling pathways mediated by importin α/β1 other than STAT1, we focused on the following signaling molecules, hypoxia inducible factor 1α (HIF-1α) and Nuclear factor-kappa B (NF-κB) component p65/RelA, since they have been shown to be transported into the nucleus by several importin α proteins ^35–37^. First, HeLa cells were transfected with AcGFP or AcGFP-ORF6, and then treated with 200 μM cobalt chloride (CoCl2) for 5 h to induce the nuclear accumulation of HIF-1α. On the other hand, the nuclear migration of NF-κB p65 was investigated using AcGFP- or AcGFP-ORF6-transfected cells treated with 20 ng/mL tumor necrosis factor-α (TNF-α) for 30 min. As a result, the nuclear accumulations of both of HIF-1α (Fig. 6F, G) and NF-κB p65 (Fig. 6H, I) were significantly suppressed in the AcGFP-ORF6-transfected cells. Notably, the inhibitory effects of ORF6 on these two importin α/β1-mediated signaling molecules were reduced compared to that on STAT1. These results totally suggest that the SARS-CoV-2 might negatively regulate the importin α/β1-mediated protein trafficking into the nucleus through the moderate binding of ORF6 to importin α1.

### Knockout of *KPNA2* enhances the replication of SARS-CoV-2

Since importin α1 might be one of the target molecules for SARS-CoV-2 ORF6, we further examined the roles of importin α1 on the propagation of SARS-CoV-2. The *KPNA2* gene, which encodes importin α1, was knocked out in Huh7-ACE2 cells using a single-guide RNA (sgKPNA2), and then the knockout (KO) cells were infected by SARS-CoV-2. After 6, 12, and 24 h post-infection, the viral RNA levels and viral titer were measured using the culture supernatant. Both viral RNA levels and viral titer significantly increased in the *KPNA2* KO cells 24 h following infection (Fig. 6J, K). These results suggest that importin α1 plays a suppressive function on the SARS-CoV-2 replication and that SARS-CoV-2 ORF6 might attenuate the function of importin α1 to achieve an efficient viral propagation.

### Global analysis for *KPNA2* in different tissues and at different ages

To further understand the role of the importin α1 in the propagation of SARS-CoV-2 and ultimately the COVID-19 pandemic, we assessed whether the patients’ importin α1 expression profiles are associated with COVID-19 symptoms. Using the GTEx dataset, we analyzed whether the importin α1 expression level in lung tissues and whole blood cells, with sexes as well as across age is correlated with symptoms using the European ancestry (EUR) samples. We found that the importin α1 levels significantly decreased with ages in lung tissues (Fig. 7A) and that the tendency was associated with males rather than female (Fig. 7B). On the other hand, the expression levels in whole blood increased in an age-dependent manner in both sexes (Fig. 7C, D). We also found that the *KPNA2* expression levels in the lungs were significantly lower in Asian ancestry than those in European, African, and other ancestries (Fig. S5). These data suggest that the low expression level of importin α1 in the lungs might represent a COVID-19-associated concern observed in older patients, particularly for males, and be one of the risk factors for infection of SARS-CoV-2.

**Figure 7.**
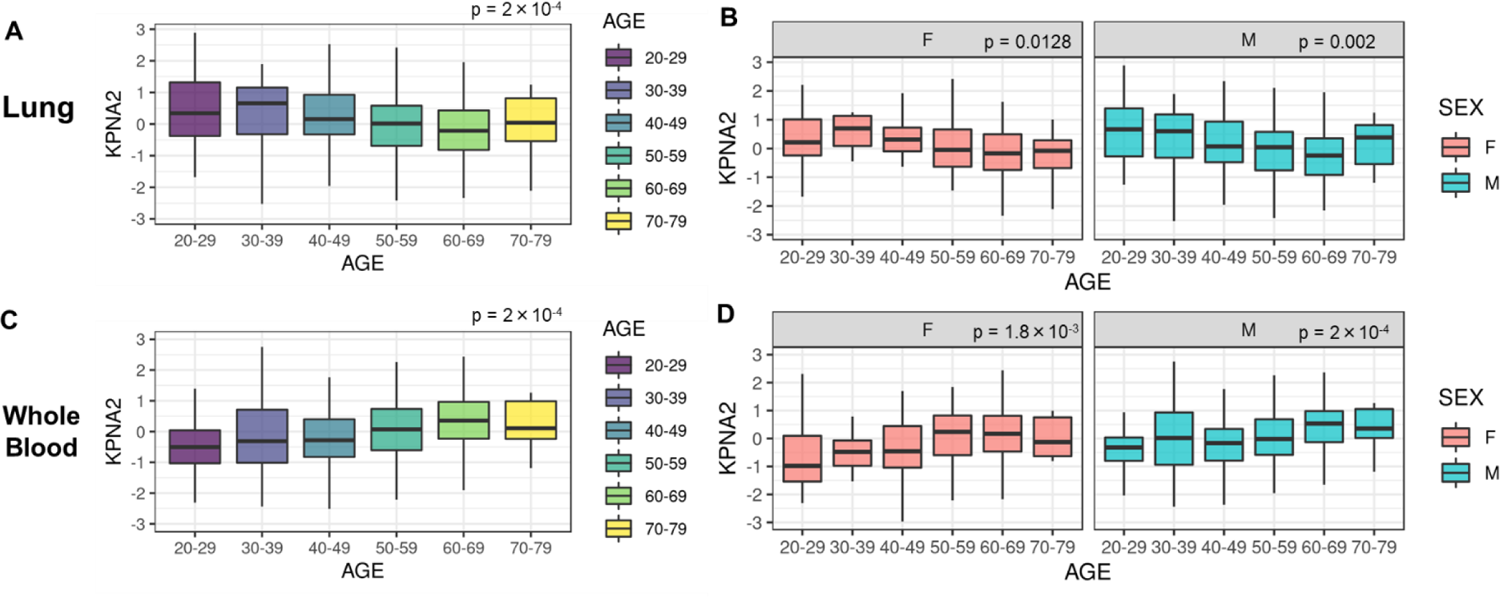
Expression levels of *KPNA2* gene for different age categories. GTEx donors whose estimated ancestry was EUR (n = 436 for lung tissues in **A** and **B**, and n = 558 for whole blood in **C** and **D**) were used. *P*-values for the trends between *KPNA2* expression levels and age categories were obtained using the two-sided Jonckheere-Terpstra test. The box represented the first and third quartiles and the center line represented the median. The upper whisker extended from the hinge to the highest value that is within the 1.5 × IQR of the hinge, the lower whisker extended from the hinge to the lowest value within the 1.5 × IQR of the hinge, and the data beyond the end of the whiskers were plotted as points. F, female; M, male; IQR, interquartile range.

## DISCUSSION

In this study, we demonstrated that SARS-CoV-2 ORF6 plays an important role in viral replication and pathogenesis of COVID-19 *in vivo*. In addition, we discovered that ORF6 acts on the nucleocytoplasmic signaling via two distinct ways. That is, first, ORF6 inhibits the nuclear translocation of one of the key signaling molecules for COVID-19, STAT1, through its direct binding to antagonize the IFN signaling. Second, ORF6 directly affects the function of importin α to impair the nuclear transportation of cNLS-containing cargos including signaling molecules such as HIF-1α and NF-κB p65.

A previous study reported that SARS-CoV ORF6 tethered importin α1/KPNA2, but not importin α5/KPNA1, to the ER and, as a result, sequestered importin β1 into the ER/Golgi segment through the interaction with importin α1, resulting in the inhibition of PY-STAT1 nuclear import ^7^. Since the nuclear transport of PY-STAT1 is known to be mediated by importin α5/KPNA1 ^24, 25^ and we showed here that importin α1 can enter the nucleus even in the presence of ORF6 under oxidative stress conditions, it is unlikely that SARS-CoV-2 ORF6 tethers importin α1 to the ER to cause the nuclear exclusion of STAT1.

In contrast, we found that the Flag-STAT1 was more abundantly localized in the cytoplasm in the absence of IFN stimulation in the ORF6-transfected cells than in the control WT cells (Fig. 4A, B). It was previously demonstrated that unphosphorylated STAT1 shuttles between the nucleus and the cytoplasm, and this shuttling might play an important role in regulating the expression of IFN stimulated genes ^12, 38–40^. Moreover, in this study, we found that the bacterially purified STAT1 proteins, which are not phosphorylated, binds to ORF6. Since the phosphorylation of STAT1 has been shown to be unaffected by ORF6 upon the IFN stimulation ^8, 10^, the interaction between ORF6 and STAT1 might occur in a phosphorylation-independent manner. Collectively, we propose a scenario that the nuclear exclusion of STAT1 is caused by the direct binding with ORF6 independently of importin α proteins.

On the other hands, we found that the subcellular localizations of all importin α subtypes were altered in ORF6-transfected cells, suggesting a possibility that ORF6 directly or indirectly influences the importin α/β1-mediated nuclear transport pathways. Consistent with this possibility, we further found that the nuclear accumulation of the mCherry-NLS substrate was significantly reduced in the AcGFP-ORF6-transfected cells. In addition, using the semi-intact nuclear transport assay, we demonstrated that the addition of ORF6 inhibits the nuclear transportation of GST-NLS-mRFP. Furthermore, we found that ORF6 negatively regulates the nuclear import of HIF-1α and NF-κB p65, which have been already shown to be mediated by importin α proteins ^35–37^. Since the nucleocytoplasmic trafficking is vital for cell survival, SARS-CoV-2, therefore, should avoid antiviral immune signaling without affecting cell survival to develop COVID-19.

Recently, it has been shown that the specific interaction of ORF6 with the NPC components, Nup98 and Rae1, might disrupt the nuclear transport ^9, 28, 41^. Consistently, it has been already known that the hijacking of the NPC components suppresses the host mRNA export system ^41^. Moreover, Miorin et al. provided a supporting evidence that the arginine substitution at residue 58 of ORF6 abolishes its IFN antagonistic function^9^. In this study, we demonstrated that STAT1 binds to ORF6 via the residues 49-61 from the C-terminus (M0) which contains a methionine amino acid at the 58 position. Furthermore, we found that this C-terminal region of ORF6 plays a critical role in altering the subcellular localization of importin α1. Taken together, we propose that the ORF6 negatively regulates the nucleocytoplasmic trafficking through the binding with importin α1 and some NPC components. Considering that various mutations and deletions have been found in ORF6 ^31, 42^, further studies will be required to understand how ORF6, especially its C-terminus region functions for the viral infection in order to utilize it as a therapeutic target.

Consistently, we found that the viral propagation of SARS-CoV-2 is enhanced in *KPNA2* KO cells. In addition, the GTEx dataset revealed that the *KPNA2* levels in lung tissue significantly decrease with the advancement in age, and that this decrease is more remarkable in males than in females. Moreover, the another GTEx analysis indicated that the levels of *KPNA2* in the lungs tend to be lower in Asians than in other genetic ancestries, although further data need to be required to make a definite conclusion, since the sample size for the Asian group is smaller than that of the other groups. Since hypoxia has been recognized as a pathogenic factor in COVID-19 patients ^43, 44^, we suspect that ORF6 might impair the nuclear import of HIF-1α, which is required for the response to hypoxia, to affect lung cell functions in the COVID-19 patients, so that the lung oxygen levels cannot be maintained appropriately. Thus, the downregulation of importin α1 may accelerate the replication of the virus in the lungs, in particular in older individuals. In conclusion, we propose that in the lungs of older individuals, SARS-CoV-2 ORF6 exhibits dual effects on the viral proliferation by inhibiting the STAT1 signaling and negatively regulating the importin α/β1-mediated nuclear transport pathways to avoid the interferon immune responses. Understanding the effects of SARS-CoV-2 proteins on the nucleocytoplasmic trafficking system might provide a novel approach for COVID-19 therapeutics.

## MATERIALS AND METHODS

### Animal care, and the production of monoclonal antibody

All animal experiments using the SARS-CoV-2 virus were performed in biosafety level 3 (ABSL3) facilities at the Research Institute for Microbial Diseases, Osaka University. The animal experiments, and the study protocol were approved by the Institutional Committee of Laboratory Animal Experimentation of the Research Institute for Microbial Diseases, Osaka University (R02-08-0). Throughout the study, we focused on minimizing animal suffering and to reducing the number of animals used in the experiments. Four weeks-old male Syrian hamsters were purchased from SLC (Shizuoka, Japan).

Experimental procedures in production of monoclonal antibody were approved by the CEC Animal Care and Use Committee (permission number: CMJ-044) and performed according to CEC Animal Experimentation Regulations. A rat monoclonal antibody that specifically recognized the SARS-CoV-2 ORF6 protein was generated using the rat medial iliac lymph node method ^45^. An 8-week-old female WKY rat was injected into the rear footpads with 100 μL of emulsions containing ORF6 peptide (CEEQPMEID)-conjugated KLH and Freund’s complete adjuvant. Seventeen days after the first immunization, an additional immunization of SARS-CoV-2 ORF6 peptide-KLH was administered without an adjuvant into the tail base of the rat. Four days after the second immunization, cells from the iliac lymph nodes of the immunized rat were fused with mouse myeloma Sp2/0-Ag14 cells at a ratio of 5:1 in 50% polyethylene glycol. The resulting hybridoma cells were plated onto 96-well plates and cultured in HAT selection medium (Hybridoma-SFM [Life Technologies, Grand Island, CA, USA]; 10% FBS; 1 ng/mL mouse IL-6; 100 μM hypoxanthine [Sigma-Aldrich, St. Louis, MO, USA]; 0.4 μM aminopterin [Sigma-Aldrich]; and 16 μM thymidine [WAKO, Osaka, Japan]). The SARS-CoV-2 ORF6-specific antibody was screened using ELISA, western blotting, and immunostaining of hybridoma supernatants. Finally, hybridoma clone producing the monoclonal antibody later named 8B10, was selected. Using a rat isotyping kit the MAb 8B10 was found to be an IgG 1 (k) antibody subtype. The monoclonal antibody against SARS-CoV-2 NP (3A9 clone) was generated by Cell Engineering Corporation (Osaka, Japan).

### Viruses

The SARS-CoV-2 (2019-nCoV/Japan/TY/WK-521/2020) strain was isolated at the National Institute of Infectious Diseases (NIID). The Germany/BavPat1/2020, USA-CA2/20200 (USA-CA2), NY-PV08410/2020, HK/VM20001061, NY-PV08449/2020, NY-PV09197/2020 were obtained from BEI Resources (Manassas, VA, USA). The different strains of SARS-CoV-2 were used to infect VeroE6/TMPRSS2 cells cultured at 37 °C with 5% CO2 in DMEM (WAKO, Osaka, Japan) containing 10% fetal bovine serum (FBS; Gibco, Grand Island, NY, USA) and penicillin/streptomycin (100 U/mL, Invitrogen, Carlsbad, CA, USA). The viral stock was generated by infecting VeroE6/TMPRSS2 cells at an MOI of 0.1. The viral supernatant was harvested at two days post infection and the viral titer was determined using plaque assay.

### Plasmid construction for mammalian expression

The AcGFP and HA were amplified and cloned into pCAGGS vector designed as pCAG AcGFP-HA. The cDNA of ORF6 was obtained from Vero-TMPRSS2 cells infected with SARS-CoV-2. The wild type ORF6, ORF6-M1, ORF6-M2 and ORF6-M3 were amplified and cloned into pCAG AcGFP-HA designed as pCAG AcGFP-ORF6-HA, pCAG AcGFP-ORF6M1-HA, pCAG AcGFP-ORF6M2-HA, and pCAG AcGFP-ORF6M3-HA, respectively. The ORF6Δ9 was constructed using the method of splicing by overlap extension method (ORF6Δ9-N and -C). The primers used throughout the study are described in Table S1. All cDNAs were amplified using polymerase chain reaction (PCR) and the Tks Gflex DNA Polymerase (Takara Bio., Shiga, Japan). The amplified cDNAs were cloned into the indicated plasmids using an In-Fusion HD cloning kit (Clontech, Mountain View, CA, USA). The sequences of all plasmids were confirmed by Eurofins Genomics (Tokyo, Japan).

Full-length STAT1 was amplified from a previously subcloned plasmid ^46^ using the primers described in Table S1. The PCR products were cloned into a pcDNA5/FRT/3xFLAG expression vector ^47^. Human importin αs including importin α1/KPNA2 and importin α5/KPNA1 were cloned into a pcDNA5/FRT/FLAG expression vector, as previously described ^47^. For constructs encoding the SV40T antigen NLS (NLS; PKKKRKVED), the relevant oligonucleotides (Table S1) were ligated into the pmCherry-C1 vector (Clontech).

### Plasmid constructions for bacterially expressed recombinant proteins

Full-length ORF6 cDNA was amplified using the specific primers described in Table S1. The PCR products were cloned into a pGEX6P2 vector (Clontech) which was subcloned the GFP gene at the N-terminus ^48^. Construct integrity was confirmed by DNA sequencing. For constructs encoding the C-terminus of ORF6 (M0), the relevant oligonucleotides (Table S1) were ligated into a pGEX2T-GFP vector, which contained the GFP gene at the multicloning site; thus, producing the pGEX2T-M0-GFP vector. The plasmid pGEX6P2/hSTAT1 was subcloned from the pcDNA5/FRT/3xFLAG expression vector. The plasmids pGEX6P3/flag-human-importin α1 and pGEX6P3/flag-human-importin α5 were obtained as previously described ^24, 47^. The relevant oligonucleotides of SV40T NLS was ligated to the pGEX2T vector containing the monomeric RFP (mRFP) gene at the multicloning site; thus, producing the pGEX2T-NLS-mRFP vector.

### Purification of bacterially expressed recombinant proteins

Purification of bacterially expressed recombinant proteins was performed as previously described ^47, 49^. Cleavage of the GST tag to induce cleaved fusion proteins was performed using PreScission protease (10 U/mg of fusion protein, GE Healthcare, Uppsala, Sweden) or thrombin protease (10 U/mg of fusion protein, Sigma-Aldrich, Germany). Importin β1, p10/NTF2, and GDP-bound Ran were purified as previously described ^47, 49^.

### Antibodies

The following primary antibodies were used in this studies: Phospho-STAT1 (Tyr701) (#9167 [58D6], Cell Signaling Technology (CST) Inc., Danvers, MA, USA), STAT1 (#9172, CST), importin α1/KPNA2 (ab84440, Abcam, Cambridge, MA, USA), HIF-1α (ab51608 [EP1215Y], Abcam), NF-κB p65 (#8242 [D14E12], CST), importin β1 (ab2811 [3E9], Abcam), CAS (ab96755, Abcam), Flag (M2 [F1804], Sigma-Aldrich), GFP (A-11122, rabbit, Thermo Fisher Scientific, Waltham, MA, USA), GFP (M048-3, mouse, MBL, Nagoya, Japan), NP (3A9, mouse mAb, Cell Engineering Co., Osaka, Japan), Actin (A2228, Sigma-Aldrich), and HA (MMS-101R, Biolegend, San Diego, CA, USA).

Horseradish peroxidase (HRP)-conjugated anti-rabbit (#111-035-003), anti-mouse (#115-035-003), or anti-rat (#112-035-003) secondary antibodies (Jackson ImmunoResearch Inc. West Grove, PA, USA) were used for western blotting. The secondary antibodies used for indirect immunofluorescence were as follows: Alexa Fluor Plus 488 conjugated anti-rabbit (A32731) or anti-mouse (A32723), and Alexa Fluor 594 conjugated anti-rabbit (A21207) or anti-mouse (A21203) (Invitrogen).

### Cell culture and transfection

HeLa cells or HEK293 cells were cultured in Dulbecco’s modified Eagle’s medium (DMEM; Invitrogen), containing 10% FBS (#10270, Gibco) at 37 °C in 5% CO_2_. The cells were plated onto 18 × 18 mm coverslips (Menzel-Glaser, Braunschweig, Germany) in 35-mm dishes for immunofluorescence or 60-mm dishes (IWAKI, Tokyo, Japan) for qRT-PCR 2 days prior to transfection. The transfections were performed using Lipofectamine 2000 DNA Transfection Reagent (Thermo Fisher Scientific) or the TransIT-LT1 Transfection Reagent (Mirus, Madison, WI, USA) following manufacturer’s instructions.

### Indirect immunofluorescence

HeLa cells were cultured on 18 × 18 mm coverslips (Matsunami, Osaka, Japan) in 35-mm dishes (IWAKI) and incubated for 48 h at 37 °C in 5% CO_2_. The reagents used for indirect immunofluorescence were as follows; IFN-β and IFN-γ (final conc. was 50 ng/mL for 30 min; Miltenyi Biotec, Bergisch Gladbach, Germany), TNF-α (final conc. was 20 ng/mL for 30 min; Miltenyi Biotec), hydrogen peroxide (H_2_O_2_, final conc. was 200 μM for 1 h), and Cobalt(II) chloride hexahydrate (CoCl2, final conc. was 200 μM for 5 h; C8661, Sigma-Aldrich). Following fixation with 3.7% formaldehyde in PBS for 15 min, cells were treated with 0.1% Triton X-100 in PBS for 5 min and then blocked in PBS containing 3% skim milk for 30 min. For the anti-Phospho-STAT1 antibody (58D6, Rabbit mAb, #9167, CST), cells were permeabilized with 100% methanol at −20 °C for 20 min and then blocked in 3% skim milk in PBS. Cells were incubated with primary antibodies (1:200) with 3% skim milk in PBS overnight at 4 °C. The following day, the cells were incubated with the Alexa-Fluor-488 plus- or Alexa-Fluor-594-conjugated secondary antibodies (Invitrogen). Nuclei were counterstained with DAPI (1:5,000 in PBS, Dojindo Laboratories, Kumamoto, Japan) for 20 min at 25 °C. The coverslips with fixed cells were mounted on glass slides using ProLong Gold Antifade (#36930, Invitrogen). Cells were examined under a confocal microscope (Leica TCS SP8 II; Leica Microsystems, Wetzlar, Germany).

### Western blotting

Western blotting was performed as previously described ^50^. The membranes were incubated with primary antibodies (dilutions ranging from 1:1000 to 1:2000) diluted in Can Get Signal Immunoreaction Enhancer Solution 1 (TOYOBO, Osaka, Japan) overnight at 4 °C. The used HRP-conjugated secondary antibodies (dilutions ranging from 1:2000 for mammalian expression to 1:10,000 for bacterially purified recombinant proteins) were diluted in Can Get Signal Immunoreaction Enhancer Solution 2 (TOYOBO) at 25 °C for 1 h.

### RNA purification and quantitative RT-PCR (qRT-PCR)

For IP-10, total RNA was isolated using ReliaPrep™ RNA Tissue Miniprep System (Promega, Madison, WI, USA) according to the manufacturer’s instructions. One microgram of total RNA and the PrimeScript RT reagent kit (Takara Bio.) were used to perform the first-strand cDNA synthesis. The PCR reaction was performed as previously described ^50^. The PCR primers including those of β-actin are described in Table S2.

For detection of N2 in SARS-CoV-2, total RNA of Huh7-ACE2 or lung homogenates were isolated using ISOGENE II (Nippon Gene, Toyama, Japan). Real-time RT-PCR was performed with the Power SYBR Green RNA-to-CT 1-Step Kit (Applied Biosystems, Foster City, CA, USA) using an AriaMx Real-Time PCR system (Agilent, Santa Clara, CA, USA). The relative quantification of the target mRNA levels was performed using the 2^-ΔΔCT^ method. β-actin was used as the housekeeping gene. The primers used are described in Table S2.

### Quantitative RT-PCR of viral RNA in the supernatant

The amount of RNA copies in the culture medium was determined using a qRT-PCR assay as previously described with slight modifications ^51^. Briefly, 5 μL of culture supernatants were mixed with 5 μL of 2× RNA lysis buffer (2% Triton X-100, 50 mM KCl, 100 mM Tris-HCl [pH 7.4], 40% glycerol, 0.4 U/μL of Superase•IN [Thermo Fisher Scientific]) and incubated at 25 °C for 10 min. Next, 90 μL of RNase free water were added to the mix. A volume of 2.5 μL of the diluted sample was added to 17.5 μL of reaction mix. Real-time RT-PCR was performed using the Power SYBR Green RNA-to-CT 1-Step Kit (Applied Biosystems) and an AriaMx Real-Time PCR system (Agilent).

### Plaque formation assay

Vero-TMPRSS2 were seeded into 24-well plates (80,000 cells/well) at 37 °C in 5% CO_2_ for overnight. The supernatants were serially diluted using inoculated medium and incubated for 2 h. Next, the culture medium was removed, fresh medium containing 1% methylcellulose (1.5 mL) was added, and the cells were cultured for 3 more days. Lastly, the cells were fixed with 4% paraformaldehyde in PBS (Nacalai Tesque, Kyoto, Japan) and the plaques were visualized by using a Giemsa’s azur-eosin-methylene blue solution (#109204, Merck Millipore, Darmstadt, Germany).

### Syrian hamster model of SARS-CoV-2 infection

Syrian hamsters were anaesthetized with isoflurane and challenged with 1.0 x 10^6^ PFU (in 60 μL) SARS-CoV-2 via intranasal routes. The body weight was monitored daily for 5 days. Five days post infection, all animals were euthanized, and the lungs were collected for histopathological examinations and qRT-PCR.

### Immunohistochemistry

Lung tissues were fixed with 10% neutral buffered formalin and embedded in paraffin. For immunohistochemical staining, 2 μm thick sections were immersed in citrate buffer (pH 6.0) and heated for 20 min with a pressure cooker. Endogenous peroxidase was inactivated by immersion in 3% H_2_O_2_ in PBS. After treatment with 5% skim milk in PBS for 30 min at 25 °C, the sections were incubated with mouse anti-NP antibody (1:500, clone 3A9). EnVision^+^ system-HRP-labeled polymer anti-mouse secondary antibody (Dako, Carpinteria, CA, USA) was used. Lastly, the sections were counterstained with hematoxylin and the positive signals were visualized using the peroxidase–diaminobenzidine reaction.

### KPNA2 knockout

For the single-guide RNA (sgRNA) targeting *KPNA2*, the targeting sequences were designed using three different sequences for each gene as previously described ^52^. The targeting sequences were synthesized using DNA oligos (Eurofins Genomics, Tokyo, Japan), and cloned into the lentiCRISPR v2 (Addgene, #52961) digested by BmsBI (New England Biolab, MA, USA). The target sequences for *KPNA2* were described in Table S3. Lentiviruses expressing three types of target sequences per gene were mixed, introduced into the Huh7 cells expressing the ACE2 receptor (Huh7-ACE2), and maintained in a culture medium supplemented with 1 µg/mL puromycin for 3 weeks. For viral infection, sgControl (sgCtl) or sgKPNA2 Huh7-ACE2 cells were seeded into 24-well plates and incubated at 37 °C for 24 h. The different SARS-CoV-2 strains were used to infect the cells (MOI 0.1) and supernatants were collected at 0, 6, 12, and 24 h. The intracellular viral RNA was quantified using qRT-PCR while the viral titers were quantified using the plaque forming assay.

### Construction of SARS-CoV-2 replicon DNA

SARS-CoV-2 replicon vector, pBAC-SCoV2-Rep, was generated using the CPER reaction as previously described ^53^, with some modifications. Briefly, seven DNA fragments covering the SARS-CoV-2 genome (excluding the region spanning from S gene to ORF8 gene) were amplified using PCR, and subcloned into a pCR-Blunt vector (Invitrogen). The DNA fragments containing cytomegalovirus (CMV) promoter, a 25 nucleotide synthetic poly(A), a hepatitis delta ribozyme as well as a bovine growth hormone (BGH) termination, and a polyadenylation sequences (the lightly shaded region in Fig. 3D) were amplified using a conventional overlap extension PCR, and subcloned into the NotI sites of pSMART BAC vector (Lucigen, Middelton, WI, USA). The luciferase reporter vector pGL4 was used as the template for PCR amplification of Renilla luciferase gene. For CPER reaction, nine DNA fragments that contain approximately 40-bp overlapping ends for two neighboring fragments were amplified by PCR using the aforementioned plasmids. Next, the PCR fragments were mixed equimolarly (0.1 pmoL each) and subjected to CPER reaction using the PrimeSTAR GXL DNA polymerase (Takara Bio.). The CPER product was extracted using phenol-chloroform, followed by ethanol precipitation, resolved in TE buffer, and transformed into the BAC-Optimized Replicator v2.0 Electrocompetent Cells (Lucigen). The replicon vector was maxipreped using a NucleoBond Xtra BAC kit (Takara Bio.).

### Generation of SARS-CoV-2 recombinant virus

SARS-CoV-2 recombinants were generated by CPER reaction as previously described ^54^ with some modifications. Briefly, 14 SARS-CoV-2 (2019-nCoV/Japan/TY/WK-521/2020) cDNA fragments (#1-#13) were amplified using PCR and subcloned into a pBlueScript KS(+) vector. The primers used are described in Table S4. The DNA fragments containing CMV promoter, a 25-nucleotide synthetic poly(A), hepatitis delta ribozyme and BGH termination and, polyadenylation sequences (#14) were synthesized by Integrated DNA Technologies (Coralville, IA, USA), and subcloned into a pBlueScript KS(+) vector. To generate a reporter SARS-CoV-2 virus, we inserted a NanoLuc (NLuc) gene and 2A peptide into the ORF6 sequence of fragment #12 (SARS-CoV-2/NLuc2AORF6). To generate an ORF6 deficient SARS-CoV-2 virus, ORF6 gene was replaced with an NLuc gene (SARS-CoV-2/ΔORF6). For CPER reaction, 14 DNA fragments that contain approximately 40- to 60-bp overlapping ends for two neighboring fragments were amplified using PCR from the subcloned plasmids. Next, the PCR fragments were mixed equimolarly (0.1 pmoL each) and subjected to CPER reaction using the PrimeSTAR GXL DNA polymerase (Takara Bio.). The cycling condition used included an initial 2 min of denaturation at 98 °C; 35 cycles of 10 s at 98 °C, 15 s at 55 °C, and 15 min at 68 °C; and a final elongation period of 15 min at 68 °C. The half of CPER product was transfected into IFNAR1-deficient HEK293 cells TransIT-LT1 transfection reagents (Mirus), according to the manufacturer’s instructions. ACE2 and TMPRSS2 receptors were induced in HEK293-3P6C33 cells using tetracycline. At 24 h post-transfection, the culture medium were replaced with DMEM containing 2% FBS and doxycycline hydrochloride (1 μg/ml). At 7-10 days post transfection, the culture medium containing progeny viruses (P0 virus) were passaged and amplified in VeroE6/TMPRSS2 cells.

### Luciferase assay

Huh7 cells were seeded into a 24-well plate and incubated at 37 °C for 24 h. The cells were transfected with pISRE-TA-Luc, pRL-TK (Promega,), and pCAG AcGFP-HA or pCAG AcGFP-ORF6-HA using TransIT-LT1 reagents (Mirus) according to the manufacturer’s instructions. The cells were incubated for 24 h after transfection and treated with IFN-γ (50 ng/mL) for 12 h.

VeroE6/TMPRSS2 cells were seeded into a 24-well plate and incubated at 37 °C for 24 h. The cells were transfected with pISRE-TA-Luc and pRL-TK (Promega). After 24 h, the cells were infected with SARS-CoV-2 and treated with IFN-γ (50 ng/mL) for 12 h. The luciferase activity was detected using the Dual-Luciferase Reporter Assay System (Promega) according to the manufacturer’s instructions.

### Semi-intact nuclear transport assay

A digitonin-permeabilized *in vitro* nuclear transport assay was performed as previously described ^47^. The NLS substrate GST-NLS-GFP was used 4 pmoL in 10 μL of reaction mixture, and the competitive substrate AcGFP-ORF6 was added to the assay with 20 pmoL, 40 pmoL, and 80 pmoL which represented 5×, 10×, or 20× the NLS-substrate dosage, respectively.

### The analyses of GTEx datasets

We obtained publicly available data regarding the expression levels of *KPNA* genes, sex, age category (20-29, 30-39, 40-49, 50-59, 60-69, 70-79), and genotype principal components (PCs) of 49 tissues from 838 post-mortem donors from GTEx v8 website (https://www.gtexportal.org/home/). These samples were used for expression quantitative trait locus (eQTL) analysis, and the expression levels were already normalized (see “3.4.2 Gene expression quantification” in the supplementary materials ^55^). From the genotype PC1 and PC2, we inferred the ancestry of the individuals (European (EUR), African (AFR), Asian, and others) by comparing with plots showing those PCs and reported the different races in the GTEx v8 paper ^56^ (Fig. S4). We performed the Jonckheere-Terpstra test (the number of permutation iterations was 10^4^) to analyze the trends between age category and *KPNA2* expression levels using R package *clinfun* v1.0.15, and the Welch t-test for quantifying the differences in expression levels between males and females.

### Statistical Analysis

Data were analyzed with Prism 7.0 software (GraphPad Software, La Jolla, CA), and expressed as the mean ± standard deviation (SD). Statistical significance was evaluated by one-way ANOVA or two-way ANOVA for comparison of multiple groups and the Student t test for two groups, *P < 0.05, **P < 0.01, ***P < 0.001.

## ACKNOWLEDGMENTS

This work was funded by the Japan Agency for Medical Research and Development (AMED) [grant numbers 20fk0108263h0001 and 20fk0108296s0101] to YM, TS, TT, and TO.

## AUTHOR CONTRIBUTION

Conceptualization: Y.M. and T.O.

Methodology: Y.M., T.S., T.T., Y.S., M.K., T.T., Y.K., Y.Y., and T.O.

Investigation: Y.M., Y.I., T.S., T.T., Y.S., M.K., C.H., C.W., M.O., and T.O.

Resources: Y.M., T.S., T.T., Y.S., K.M., T.T., Y.K., T.O., and M.O.

Writing – Original Draft: Y.M., Y.I., T.S., T.T., Y.S., M.K., T.T., Y.K., Y.Y., T.O., and M.O.

Writing – Review & Editing: Y.M., Y.I., T.S., T.T., Y.S., M.K., C.H., C.W., M.O., K.M., T.T., Y.K., Y.Y., T.O., and M.O.

Funding Acquisition: Y.M., T.S., T.T., and T.O Supervision: Y.M. and T.O.

