## Supplemental Figures for "SARS-CoV-2 ORF6 disturbs nucleocytoplasmic trafficking to advance the viral replication"

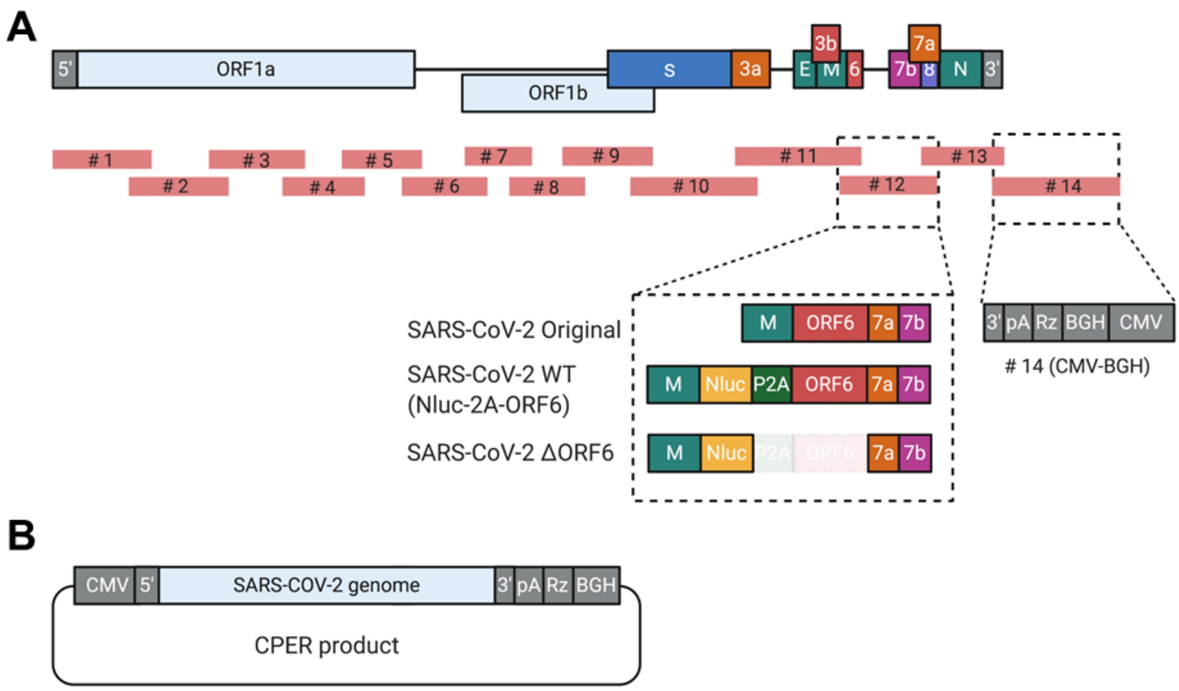


**Figure S1. Establishment of recombinant SARS-CoV-2 by the circular polymerase extension reaction**

**A.** Schematic representation of recombinant SARS-CoV-2. A total of 13 fragments (#1-#13) that cover the viral genome were amplified with 40-60-nt overlapping ends. The fragments were mixed with the #14 fragment, which contained the CMV promoter (CMV) followed by the first 20 nt of SARS-CoV-2 genome and, at the other end, the 3' UTR of SARS-CoV-2, a synthetic poly(A) tail (pA), hepatitis delta virus ribozyme (Rz), and bovine growth hormone polyadenylation sequence (BGH). NLuc, NanoLuc gene; P2A, Porcine teschovirus 2A peptide. **B.** The genetic structure of the CPER product of the recombinant SARS-CoV-2. A total of 14 fragments were assembled by CPER.

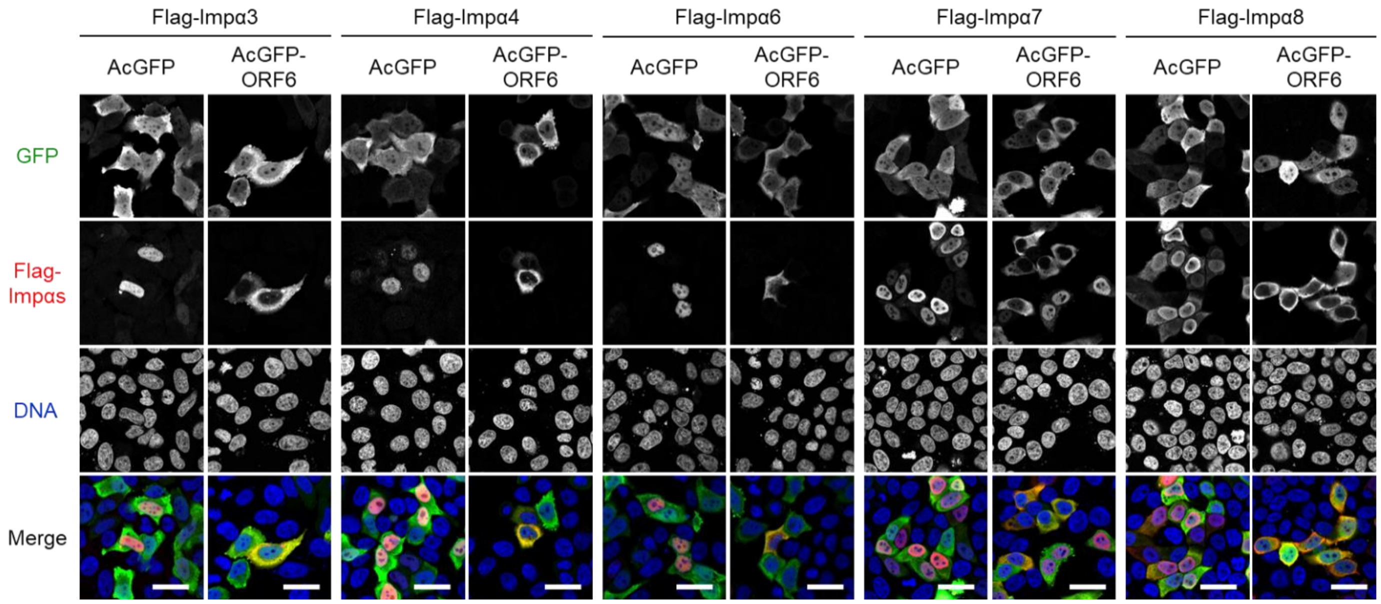


**Figure S2. Subcellular distribution of Flag-importin α subtypes**

Subcellular localization of Flag-importin α3, α4, α6, α7, and α8 in HeLa cells expressed with either AcGFP or AcGFP-ORF6. The Flag-importin α proteins and the GFP protein were detected using the anti-Flag antibody or anti-GFP antibody, respectively. DNA was stained with DAPI. Scale bars: 30 μm.

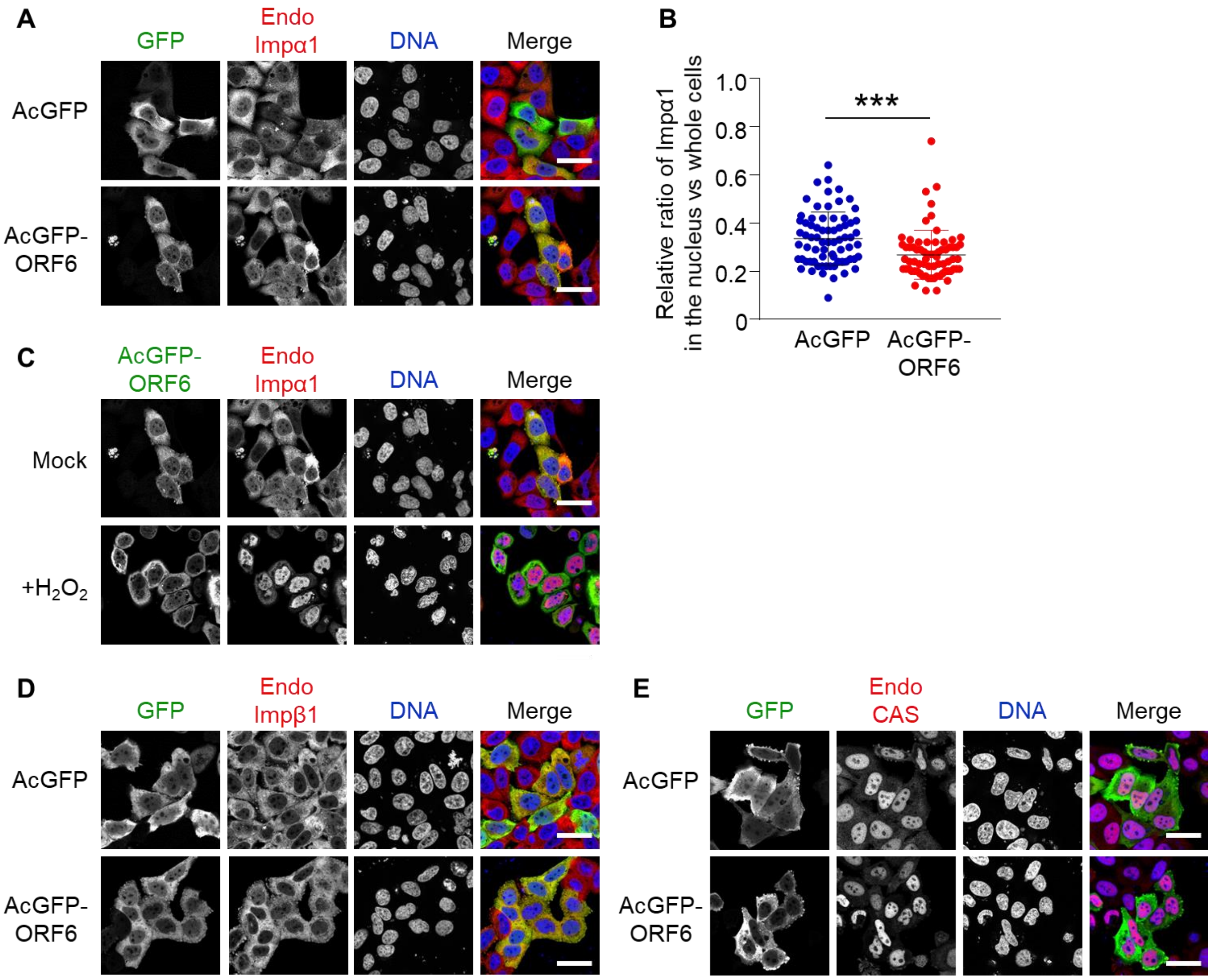


**Figure S3. Importin α1 shuttles in ORF6 expressed cells**

**A.** Subcellular localization of endogenous importin α1 (Endo Impα1) in HeLa cells expressed with either AcGFP or AcGFP-ORF6. The importin α1 proteins and the GFP protein were detected using the anti-importin α1 antibody or anti-GFP antibody, respectively. DNA was stained with DAPI. Scale bars: 30 µm. **B.** The graph represents the relative values of the fluorescent in the nucleus against the whole cells in A. Signal intensities of total 70 different nuclei from two independent experiments were measured, and statistically analyzed using student's t-test (***$P<0.001$, error bars represent SD). **C.** Subcellular localization of endogenous importin α1 (Endo Impα1) in HeLa cells expressed with AcGFP-ORF6 with or without hydrogen peroxide (200 µM $H_2O_2$) for 30 min. The importin α1 proteins and the GFP protein were detected using the anti-importin α1 antibody or anti-GFP antibody, respectively. DNA was stained with DAPI. Scale bars: 30 µm. **D-E.** Subcellular localization of endogenous importin β1 (Endo Impβ1) or CAS co-transfected with either AcGFP or AcGFP-ORF6 in HeLa cells. Importin β1, CAS and the GFP protein were detected using specific antibodies. DNA was stained with DAPI. Scale bars: 30 µm.

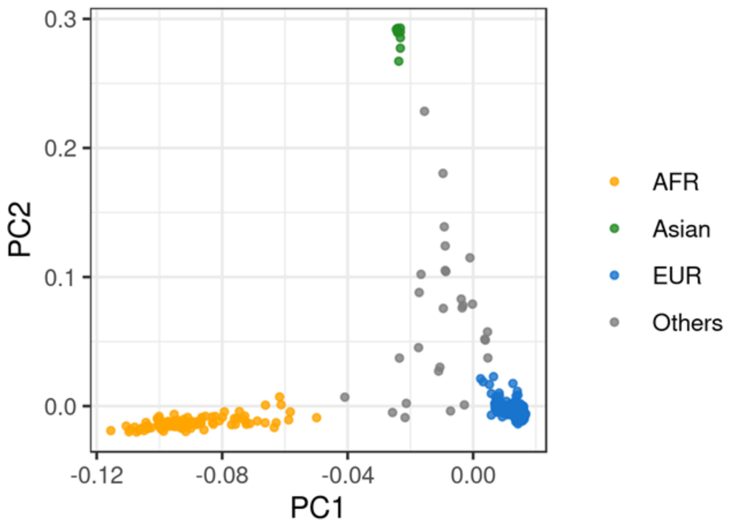


**Figure S4. Top two genotype PCs and inferred ancestry of GTEx donors**

838 GTEx donors for eQTL analysis in the GTEx v8 paper were analyzed. EUR (n=698), AFR (n=103), Asian (n=9), and others (n=28) were inferred ancestry, defined in the current study.

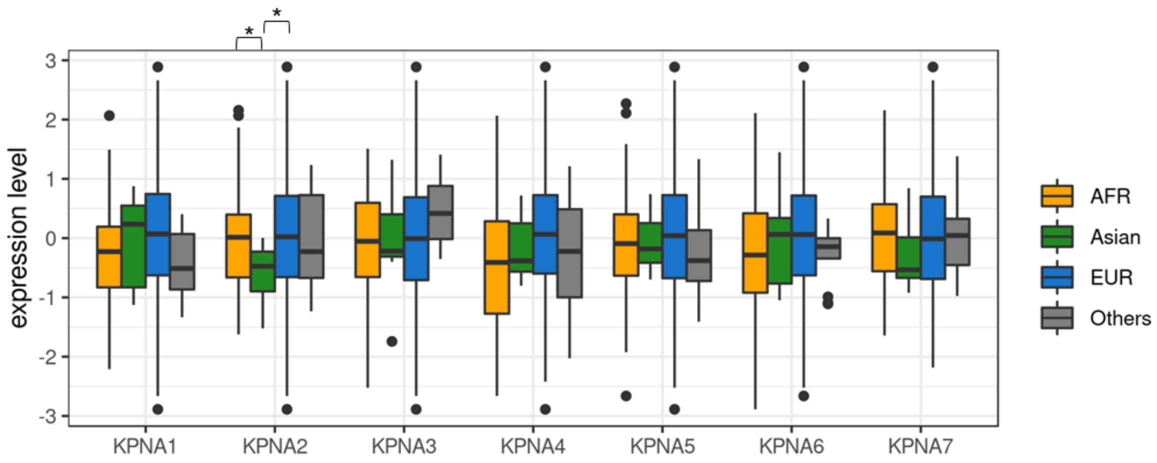


**Figure S5. Distribution of expression levels of KPNA genes in lung tissue.**

GTEx donors whose estimated ancestry was EUR (n=436), AFR (n=58), Asian (n=7), and others (n=14) were used. *, $p < 0.05$ (two-tailed Welch t-test). The box represented the first and third quartiles, the center line represented the median, the upper whisker extended from the hinge to the highest value that is within 1.5×IQR (inter-quartile range) of the hinge, the lower whisker extended from the hinge to the lowest value within 1.5× IQR of the hinge, and the data beyond the end of the whiskers were plotted as points.
