## Supplemental Tables for "SARS-CoV-2 ORF6 disturbs nucleocytoplasmic trafficking to advance the viral replication"

**Table S1. Oligo sequences for plasmid constructions**

| | | | |
|---|---|---|---|
| Mammalian expression vector | ORF6_Fw | 5'-GCTGTACAAGGAATTCATGTTTCATCTCGTTGACTTTCAGG-3' | pCAG AcGFP-HA |
| | ORF6_Rv | 5'-GGTATCCTCTGAATTCATCAATCTCCATTGGTTGCTC-3' | |
| | ORF6-M1_Fw | 5'-GCTGTACAAGGAATTCATGTTTCATCTCGTTGACTTTCAGG-3' | |
| | ORF6-M1_Rv | 5'-GGTATCCTCTGAATTCATCAATCTCCATTGGTTGCTCTTCATCTGCTGCTGCTGCTTTATTCTCA GTTAGTGACT-3' | |
| | ORF6-M2_Fw | 5'-GCTGTACAAGGAATTCATGTTTCATCTCGTTGACTTTCAGG-3' | |
| | ORF6-M2_Rv | 5'-GGTATCCTCTGAATTCATCAATCTCCATTGGTTGTGCTGCTGCTAATTGAGAATATTTATTCT-3' | |
| | ORF6-M3_Fw | 5'-GCTGTACAAGGAATTCATGTTTCATCTCGTTGACTTTCAGG-3' | |
| | ORF6-M3_Rv | 5'-GGTATCCTCTGAATTCTGCTGCTGCTGCTGCCTCTTCATCTAATTGAGAAT-3' | |
| | ORF6 Δ9-N_Fw | 5'-GCTGTACAAGGAATTCATGTTTCATCTCGTTGACTTTCAGG-3' | |
| | ORF6 Δ9-N_Rv | 5'-ATGATGTAAGTCCTCATAATAATTAGTAATATC-3' | |
| | ORF6 Δ9-C_Fw | 5'-TGAGGACTTACATCATAAACCTCATAATTAAAA-3' | |
| | ORF6 Δ9-C_Rv | 5'-GGTATCCTCTGAATTCATCAATCTCCATTGGTTGCTC-3' | |
| | STAT1_Fw | 5’-CGATGACAAGGGATCCATGTCTCAGTGGTACGAAC-3’ | pcDNA5/FRT/3xFLAG |
| | STAT1_Rv | 5’-GGCTCTCGCTGAATTCCTATACTGTGTTCATCATACT-3’ | |
| | SV40T-NLS_Fw | 5'-GATCTCCTCCAAAAAAGAAGAGAAAGGTAGAAGACG-3' | mCherry-C1 |
| | SV40T-NLS_Rv | 5'-TCGACGTCTTCTACCTTTCTCTTCTTTTTTGGAGGA-3' | |
| Bacteria expression | ORF6_Fw | 5’- GACGATATCAGGATCCATGTTTCATCTCGTTGACTTTCAGG-3’ | pGEX6P2-GFP |
| | ORF6_Rv | 5’- GATGCGGCCGCTCGAGATCAATCTCCATTGGTTGCTC-3’ | |
| | M0_Fw | 5’- GATCCGAATTCTATTCTCAATTAGATGAAGAGCAACCAATGGAGATTGATCCC-3’ | pGEX2T-GFP |
| | M0_Rv | 5’-GGGATCAATCTCCATTGGTTGCTCTTCATCTAATTGAGAATAGAATTCG-3’ | |

**Table S2. Primers for qRT-PCR**

| | |
|---|---|
| IP-10_Fw | 5’-GGCCATCAAGAATTTACTGAAAGCA-3’ |
| IP-10_Rv | 5’-TCTGTGTG GTCCATCCTTGGAA-3’ |
| β-actin for IP-10_Fw | 5'-TTCCAGGAGCGAGATCCCT-3' |
| β-actin for IP-10_Rv | 5'-CACCCATGACGAACATGGG-3' |
| SARS-CoV-2_N2_Fw | 5’-AAATTTTGGGGACCAGGAAC-3’ |
| SARS-CoV-2_N2_Rv | 5’-TGGCAGCTGTGTAGGTCAAC-3’ |
| β-actin for N2_Fw | 5’-TTGCTGACAGGATGCAGAAG-3’ |
| β-actin for N2_Rv | 5’-GTACTTGCGCTCAGGAGGAG- 3’ |

**Table S3. sgRNA target sequences for *KPNA2***

| | |
|---|---|
| sgKPNA2_1 | 5’-AATGAGGCGTCGCAGAATAG-3’ |
| sgKPNA2_2 | 5’-TCTTCTAGGGCACTGTAAAT-3’, |
| sgKPNA2_3 | 5’-ATTTCGGAATCAAACCAGCC-3’ |
| sgControl | 5’-CCATATCGGGGCGAGACATG-3’ |

**Table S4. Primers for CPER reaction**

| | |
|---|---|
| CPER#1-Fw | 5’-GAGCTCGTTTAGTGAACCGTATTAAAGGTTTATACCTTCC-3’ |
| CPER#1-Rv | 5’-CTATCACAGTGTCATCACCAAAAGTAACCTTTGTTGGTGC-3’ |
| CPER#2-Fw | 5’-CCTTCACACTCAAAGGCGGTGCACCAACAAAGGTTACTTT-3’ |
| CPER#2-Rv | 5’-CTTCTACACCCTTAAGGGTTGTCTGCTGTTGTCCACAAGT-3’ |
| CPER#3-Fw | 5’-TCTTGAACGTGGTGTGTAAAACTTGTGGACAACAGCAGAC-3’ |
| CPER#3-Rv | 5’-TAACTTTAATTAACTGCTTCAACCAATTATTAACAATTTT-3’ |
| CPER#4-Fw | 5’-AGATAGCACTTAAGGGTGGTAAAATTGTTAATAATTGGTT-3’ |
| CPER#4-Rv | 5’-GAACCCTTAATAGTGAAATTGGGCCTCATAGCACATTGGT-3’ |
| CPER#5-Fw | 5’-TGGTTCACCATCTGGTGTTTACCAATGTGCTATGAGGCCC-3’ |
| CPER#5-Rv | 5’-TTTATGTCTACAGCACCCTGCATGGAAAGCAAAACAGAAA-3’ |
| CPER#6-Fw | 5’-TCACTACTTTCTGTTTTGCTTTCCATGCAGGGTGCTGTAG-3’ |
| CPER#6-Rv | 5’-ACCAGAAGCAGCGTGCATAGCAGGGTCAGCAGCATACACA-3’ |
| CPER#7-Fw | 5’-CTTAGTTTTAAGGAATTACTTGTGTATGCTGCTGACCCTG-3’ |
| CPER#7-Rv | 5’-TGGGTAAGCATCTATAGCTAAAGACACGAACCGTTCAATC-3’ |
| CPER#8-Fw | 5’-CAGATGGTACACTTATGATTGAACGGTTCGTGTCTTTAGC-3’ |
| CPER#8-Rv | 5’-AGGTGTGTAGGTGCCTGTGTAGGATGTAACCCAGTGATTA-3’ |
| CPER#9-Fw | 5’-AAGATTGTAGTAAGGTAATCACTGGGTTACATCCTACACA-3’ |
| CPER#9-Rv | 5’-GACTAGAGACTAGTGGCAATAAAACAAGAAAAACAAACAT-3’ |
| CPER#10-Fw | 5’-TGTTAACAACTAAACGAACAATGTTTGTTTTTCTTGTTTT-3’ |
| CPER#10-Rv | 5’-CAAATGAGGTCTCTAGCAGCAATATCACCAAGGCAATCAC-3’ |
| CPER#11-Fw | 5’-GCTTCATCAAACAATATGGTGATTGCCTTGGTGATATTGC-3’ |
| CPER#11-Rv | 5’-TAAGCTCTTCAACGGTAATAGTACCGTTGGAATCTGCCAT-3’ |
| CPER#12-Fw | 5’-TTGGAACTTTAATTTTAGCCATGGCAGATTCCAACGGTAC-3’ |
| CPER#12-Rv | 5’-TACATTCTTGGTGAAATGCAGCTACAGTTGTGATGATTCC-3’ |
| CPER#13-Fw | 5’-AAATTTCTTGTTTTCTTAGGAATCATCACAACTGTAGCTG-3’ |
| CPER#13-Rv | 5’-TTGTCATTCTCCTAAGAAGCTA-3’ |
| CPER#14-Fw | 5’-CTATCCCCATGTGATTTTAATAGCTTCTTAGGAGAATGAC-3’ |
| CPER#14-Rv | 5’-GGAAGGTATAAACCTTTAATACGGTTCACTAAACGAGCTC-3’ |
